# An ultrasensitive genetically encoded voltage indicator uncovers the electrical activity of non-excitable cells

**DOI:** 10.1101/2023.10.05.560122

**Authors:** Philipp Rühl, Anagha G. Nair, Namrata Gawande, Sassrika N.C.W. Dehiwalage, Lukas Münster, Roland Schönherr, Stefan H. Heinemann

## Abstract

Genetically encoded voltage indicators (GEVIs) are powerful, non-invasive tools for recording action potentials in excitable cells. However, most animal cell types are non-excitable, and yet variations in the membrane potential are biologically relevant in these cells as well. Resolving such small voltage signals demands GEVIs with exceptionally high sensitivity. In this study, we applied structure-guided engineering to the GEVI ASAP3 to generate rEstus, a sensor with optimized brightness, voltage sensitivity, and voltage range. rEstus is most sensitive in the resting voltage range of non-excitable cells, exhibits a 3.6-fold improvement in fast voltage spike detection, and allows for absolute voltage calibration at the single-cell level. Using rEstus, we resolved endogenous voltage fluctuations in several non-excitable cell types and demonstrate that correlation analysis of these optically recorded fluctuations provides an easy, non-invasive, real-time readout of electrical gap-junction coupling. Our work provides greatly enhanced tools and methods for the non-invasive study of electrical signaling in excitable and non-excitable cells.

## Introduction

Electrically excitable animal cells, such as neurons, muscle cells, and certain endocrine cells, are distinguished from non-excitable cells by their ability to generate action potentials characterized by large excursions of the electrical membrane voltage (V_m_) – a distinctive feature absent in non-excitable cells (1,2). However, it has been implicated for decades that variations in V_m_ occur during cell cycle progression (3). Moreover, long-distance V_m_ gradients are reported to play a regulatory role in tissue development and wound healing (4,5). Ion channels capable of changing V_m_, such as voltage-gated K^+^ channels, are required for the proper activation and migration of immune cells (6,7). Furthermore, certain types of voltage-gated Na^+^ and K^+^ channels are strongly upregulated in various types of cancers (8–12). Similarly, connexins, which establish direct electrical connections between adjacent cells, are frequently found to be dysregulated in cancer (13). Congenital mutations affecting various voltage-gated ion channels and connexins were reported to lead to severe dysmorphic phenotypes (14,15), suggesting physiological roles of these ion transport proteins beyond action-potential-based signaling. Despite increasing evidence highlighting the significance of electrical signaling in non-excitable tissues, the underlying molecular and cellular mechanisms are largely elusive (16). This knowledge gap is partly attributable to the lack of suitable non-invasive tools capable of accurately measuring subtle changes in V_m_ (17–19).

Genetically encoded voltage indicators (GEVIs), fluorescent membrane proteins that change their brightness upon V_m_ alteration, have emerged as powerful tools for measuring electrical signaling in excitable cells (20–26). Among the best performing GEVIs are derivatives of ASAP1 (Accelerated Sensor for Action Potentials 1), which features bright fluorescence and a particularly fast voltage response (20). ASAP1 contains transmembrane segments 1-4 (S1-S4) of the voltage-sensing domain (VSD) of the *Gallus gallus* voltage-sensitive phosphatase (VSP), fused to a circularly permuted green fluorescent protein (cpGFP), which was inserted between S3 and S4 of the VSD (20). Positively charged residues (sensing charges) in S4 respond to changes in V_m_, causing S4 to move inward (down state) at negative voltages or outward (up state) at positive voltages (27–30). This movement leads to a dimming of the coupled cpGFP when V_m_ changes in the positive direction. Several ASAP1 derivatives, such as ASAP3, postASAP, JEDI-2P, ASAP4b, and ASAP4e have been successfully applied to record neuronal and cardiac electrical signaling in human induced pluripotent stem cells or even in brains of living mice (20,23–25,31–33).

Since GEVIs are primarily developed for applications in excitable cells, crucial parameters of available GEVIs are not ideal for use in non-excitable cell types. Detecting minute changes in V_m_ requires bright GEVIs that generate a large fractional brightness change within the V_m_ range of interest. Because the brightness-voltage relationship of VSD-based GEVIs in the physiological V_m_ range is sigmoid, the V_m_ resolving power is strongly dependent on V_m_. In general, a non-linear response curve is desirable to detect small V_m_ changes because it allows for generating large brightness changes in the V_m_ range of interest (34). However, an increased steepness restricts the dynamic range to a narrowed V_m_ span. Thus, the location of mid-potential (V_half_) of the brightness–voltage relationship, which also defines the sensitivity optimum of the sensor, becomes highly relevant. In addition, measurements of slow changes in V_m_ during, for example, the cell cycle or the quantitative comparison of V_m_ in independent cell populations, require calibratable fluorescence signals that are independent of variations in the expression level (35,36). While several attempts by us and others were made to calibrate the signal of GEVIs using methods such as fluorescence lifetime imaging (37), pump-probe protocols (38), or dual-color ratiometry (35), none of these approaches achieved V_m_ calibration at the single-cell level.

The aim of this study was to develop a sensor optimized for V_m_ recordings in non-excitable cells regarding brightness, sensitivity, voltage range, and calibratability while still being fast enough for neurophysiological applications. Here, we present rEstus, a fast, ultrabright, ultrasensitive, and calibratable GEVI based on ASAP3. rEstus has sufficient resolution to unveil endogenous V_m_ fluctuations in non-excitable cells and enables high-fidelity detection of fast voltage spikes. The high resolving power of rEstus allows for the non-invasive evaluation of electrical gap-junction coupling between cells, exclusively based on the correlation of subtle endogenous V_m_ fluctuations.

## Results

### Design and characterization of an ultrabright and ultrasensitive voltage indicator

To derive the molecular brightness from the fluorescence intensity, we fused the red fluorescent protein mKate2 with ASAP3 (Fig. 1A). Because both green (F_cpGFP_) and red fluorescence signals (F_mKate2_) scale equally with the number of sensor proteins, the resulting fluorescence ratio (F_cpGFP_/F_mKate2_) is directly proportional to the molecular brightness ratio of the two proteins. To quantify the molecular brightness of ASAP3 as a percentage of the molecular brightness of EGFP (%_EGFP_), for which the absolute molecular brightness is known (39), we generated an mKate2-EGFP fusion protein and normalized F_cpGFP_/F_mKate2_ to F_EGFP_/F_mKate2_ (Fig. 1B).

**Fig. 1.**
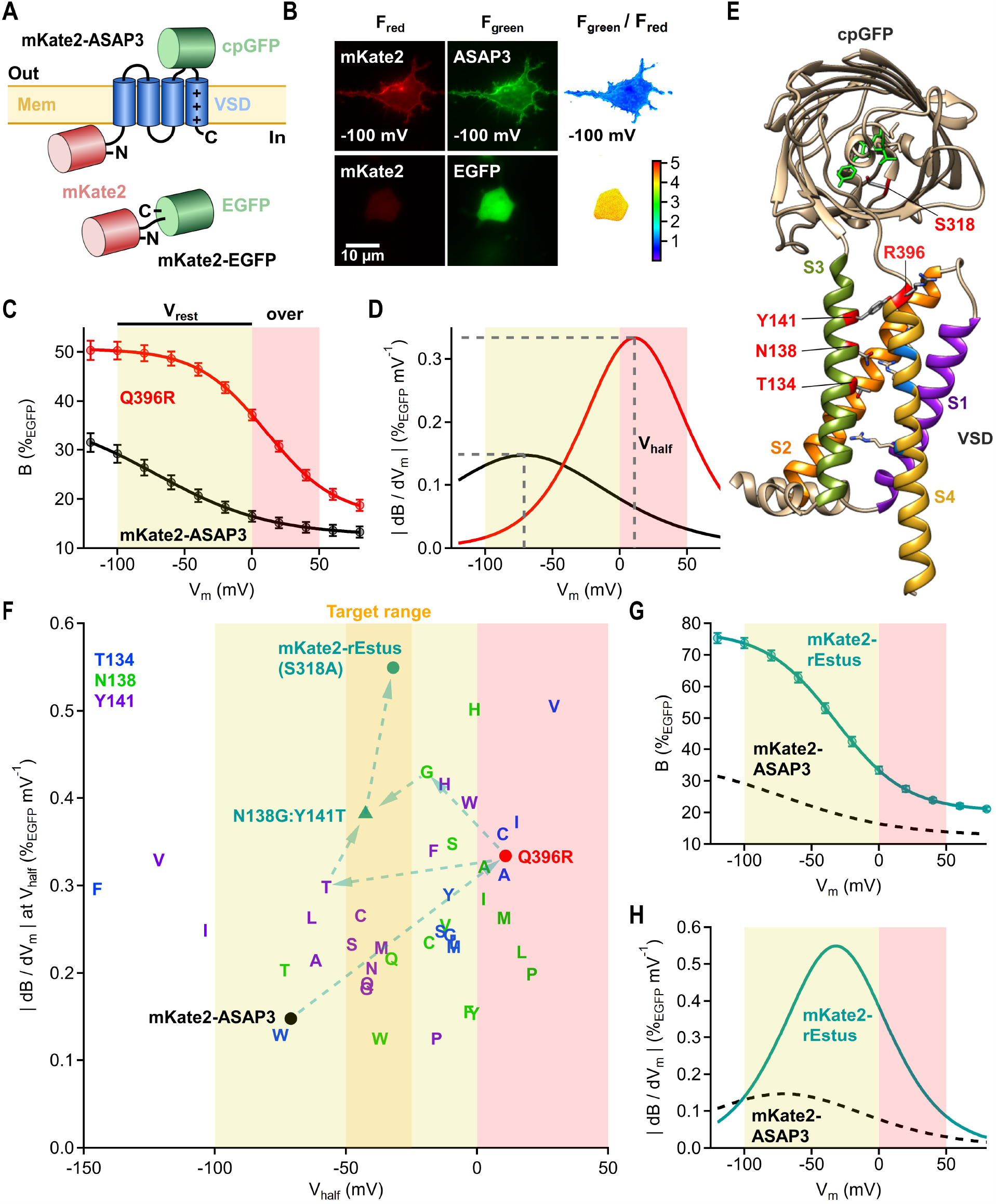
Optimization of molecular brightness, V_m_ range, and sensitivity of ASAP3. (**A**) Schematics of mKate2-ASAP3 and mKate2-EGFP. (**B**) Fluorescence and pixel-by-pixel ratio images of a HEK293T cell expressing mKate2-ASAP3 voltage-clamped at -100 mV under whole-cell patch-clamp condition (top) or expressing mKate2-EGFP without electrophysiological control (bottom). (**C**) Molecular brightness of ASAP3 and ASAP3:Q396R as a function of V_m_. F_cpGFP_/F_mKate2_ was normalized to F_EGFP_/F_mKate2_ to obtain B(%_EGFP_). Data are means ± SEM with superimposed fits (Eq. 1). (**D**) Brightness changes per voltage increment as a function of V_m_. (**E**) 3D model of ASAP3:Q396R (62). Labeled residues were mutated. (**F**) Maximum |dB/dV_m_| plotted against V_half_ for all functional constructs. T134, N138, and Y141 variants are based on mKate2-ASAP3:Q396R. Only mean values are shown, SEM values were omitted for clarity. (**G** and **H**), as in A, D but for mKate2-rEstus. The sigmoid fit for mKate2-ASAP3 (from C) is shown as black dashed lines for comparison. For n values and fit parameters refer to table 1.

We recorded the brightness–voltage relationship of ASAP3 using fluorescence imaging combined with whole-cell voltage clamp of transfected HEK293T cells. At very negative voltages, ASAP3 reached a maximum brightness (B_max_) of 37.5 ± 2.2%_EGFP_, which decreased by 24.9 ± 1.4%_EGFP_ (ΔB_max_) at saturating positive voltages (Eq. 1, Fig. 1C). However, owing to a voltage of half-maximum brightness change (V_half_) of -71 ± 3 mV and a shallow voltage dependence (k_s_) of 42.3 ± 1.1 mV, only about half of the brightness change occurred within the physiological resting V_m_ range (V_rest_) between -100 and 0 mV.

**Table 1.** Parameters of the fits according to Eq. 1 describing the brightness–voltage relationships of all constructs from Fig. 1 and Figs. S1 to S4. All constructs are based on mKate2-ASAP3.

| GEVI | *n* | B<sub>max</sub><br>(%EGFP) | SEM<br>(%EGFP) | ΔB<sub>max</sub><br>(%EGFP) | SEM<br>(%EGFP) | V<sub>half</sub><br>(mV) | SEM<br>(mV) | k<sub>s</sub><br>(mV) | SEM<br>(mV) |
| --- | --- | --- | --- | --- | --- | --- | --- | --- | --- |
| mKate2-ASAP3 | 6 | 37.5 | ± 2.2 | 24.9 | ± 1.4 | -71 | ± 3 | 42.3 | ± 1.1 |
| Q396R | 6 | 50.6 | ± 1.0 | 34.2 | ± 1.5 | 10.9 | ± 2.1 | 25.61 | ± 0.25 |
| Q396R:N138G:Y141T | 5 | 54.3 | ± 1.9 | 39.7 | ± 1.4 | -42.2 | ± 2.0 | 25.7 | ± 0.3 |
| Q396R:N138G:Y141T:S318A<br>(mKate2-rEstus) | 5 | 77.5 | ± 1.7 | 57.0 | ± 1.0 | -32.0 | ± 2.4 | 26.0 | ± 0.4 |
| Q396R:T134A | 5 | 46 | ± 6 | 32 | ± 4 | 10.3 | ± 1.9 | 25.0 | ± 0.6 |
| Q396R:T134C | 5 | 49.0 | ± 0.6 | 36.7 | ± 0.4 | 9.8 | ± 1.3 | 25.6 | ± 0.5 |
| Q396R:T134F | 5 | 61 | ± 5 | 44 | ± 5 | -146 | ± 6 | 37.2 | ± 2.3 |
| Q396R:T134G | 6 | 51 | ± 3 | 32.6 | ± 1.5 | -10.3 | ± 1.6 | 33.6 | ± 0.4 |
| Q396R:T134I | 6 | 56 | ± 3 | 37.0 | ± 1.4 | 15.1 | ± 1.1 | 25.1 | ± 0.6 |
| Q396R:T134L | 5 | 45.1 | ± 2.2 | 33.1 | ± 1.5 | -8 | ± 4 | 35.4 | ± 1.1 |
| Q396R:T134M | 5 | 54 | ± 4 | 36 | ± 3 | -8.9 | ± 1.9 | 38.6 | ± 0.8 |
| Q396R:T134S | 5 | 51 | ± 4 | 34 | ± 3 | -14.0 | ± 1.9 | 34.7 | ± 0.8 |
| Q396R:T134V | 5 | 59.1 | ± 1.8 | 44.3 | ± 1.5 | 29.4 | ± 2.2 | 22.0 | ± 0.4 |
| Q396R:T134W | 5 | 47 | ± 3 | 29.3 | ± 1.9 | -75 | ± 6 | 58 | ± 3 |
| Q396R:T134Y | 5 | 55 | ± 3 | 38 | ± 3 | -10.9 | ± 0.9 | 32.9 | ± 1.1 |
| Q396R:N138A | 5 | 45.6 | ± 2.2 | 30.5 | ± 2.0 | 2.7 | ± 2.2 | 25.0 | ± 0.7 |
| Q396R:N138C | 6 | 47.0 | ± 1.5 | 32.5 | ± 1.2 | -18.3 | ± 1.4 | 34.6 | ± 0.9 |
| Q396R:N138F | 5 | 44.2 | ± 2.4 | 21.6 | ± 1.0 | -3 | ± 3 | 34.9 | ± 1.1 |
| Q396R:N138G | 5 | 52.1 | ± 1.4 | 38.5 | ± 1.3 | -19.1 | ± 2.2 | 22.5 | ± 0.6 |
| Q396R:N138H | 5 | 58.5 | ± 2.1 | 40.7 | ± 2.0 | -1 | ± 3 | 20.3 | ± 0.3 |
| Q396R:N138I | 5 | 51.0 | ± 1.5 | 33.7 | ± 1.3 | 2.4 | ± 1.4 | 29.8 | ± 0.7 |
| Q396R:N138L | 6 | 56 | ± 3 | 38 | ± 3 | 17.0 | ± 0.9 | 42.6 | ± 1.2 |
| Q396R:N138M | 6 | 57 | ± 3 | 39 | ± 3 | 10.3 | ± 2.3 | 37.5 | ± 1.3 |
| Q396R:N138P | 6 | 41.9 | ± 0.6 | 24.1 | ± 0.8 | 21 | ± 3 | 30.5 | ± 0.8 |
| Q396R:N138Q | 5 | 55.0 | ± 0.7 | 37.0 | ± 1.2 | -33 | ± 3 | 42.8 | ± 0.37 |
| Q396R:N138S | 5 | 48.0 | ± 1.2 | 34.6 | ± 0.7 | -9.5 | ± 1.5 | 24.9 | ± 0.6 |
| Q396R:N138T | 5 | 45.2 | ± 0.9 | 23.8 | ± 1.0 | -73.3 | ± 1.5 | 29.4 | ± 0.4 |
| Q396R:N138V | 8 | 56 | ± 3 | 40.3 | ± 2.3 | -12.1 | ± 2.2 | 39.7 | ± 0.7 |
| Q396R:N138W | 6 | 46 | ± 4 | 30 | ± 4 | -37 | ± 7 | 61 | ± 7 |
| Q396R:N138Y | 6 | 47 | ± 3 | 25.8 | ± 1.8 | -1 | ± 4 | 42.1 | ± 2.0 |
| Q396R:Y141A | 5 | 47.0 | ± 2.1 | 31.7 | ± 0.5 | -61.6 | ± 2.3 | 37.1 | ± 0.7 |
| Q396R:Y141C | 5 | 45.0 | ± 2.0 | 31 | ± 3 | -44.4 | ± 1.9 | 29.4 | ± 0.7 |
| Q396R:Y141F | 5 | 58.2 | ± 2.3 | 43.4 | ± 1.7 | -17 | ± 3 | 32.0 | ± 0.5 |
| Q396R:Y141G | 6 | 45.4 | ± 2.1 | 30.1 | ± 1.3 | -42 | ± 4 | 41.5 | ± 1.4 |
| Q396R:Y141H | 6 | 63 | ± 3 | 45.0 | ± 2.0 | -12.4 | ± 0.5 | 26.99 | ± 0.24 |
| Q396R:Y141I | 6 | 45.4 | ± 2.0 | 33.2 | ± 1.8 | -103.7 | ± 1.4 | 33.5 | ± 1.0 |
| Q396R:Y141L | 5 | 41.6 | ± 2.4 | 29.3 | ± 2.0 | -63.1 | ± 1.1 | 27.9 | ± 0.3 |
| Q396R:Y141M | 5 | 46.5 | ± 1.9 | 31.2 | ± 2.2 | -37 | ± 3 | 34.2 | ± 0.9 |
| Q396R:Y141N | 5 | 47.0 | ± 2.1 | 32.1 | ± 1.9 | -40.2 | ± 1.5 | 39.2 | ± 1.0 |
| Q396R:Y141P | 5 | 46 | ± 3 | 26 | ± 3 | -15 | ± 3 | 52 | ± 5 |
| Q396R:Y141Q | 5 | 51.2 | ± 1.5 | 35.9 | ± 1.7 | -42 | ± 3 | 47.8 | ± 0.7 |
| Q396R:Y141S | 5 | 44.8 | ± 1.5 | 30.2 | ± 1.3 | -48 | ± 3 | 32.7 | ± 0.7 |
| Q396R:Y141T | 7 | 49 | ± 3 | 34.7 | ± 2.0 | -57 | ± 3 | 29.1 | ± 0.3 |
| Q396R:Y141V | 6 | 59 | ± 5 | 44 | ± 6 | -121 | ± 5 | 33.1 | ± 0.8 |
| Q396R:Y141W | 5 | 58.8 | ± 1.8 | 40.5 | ± 0.9 | -2.9 | ± 1.2 | 25.6 | ± 0.4 |
| Q396R:S318A | 5 | 80 | ± 3 | 56 | ± 3 | 21.2 | ± 2.4 | 25.6 | ± 0.7 |
| Q396R:V170T | 4 | 36 | ± 3 | 20.0 | ± 2.0 | 7 | ± 3 | 25.1 | ± 1.1 |
| Q396R:S178G | 5 | 46.6 | ± 1.7 | 33.1 | ± 1.6 | 9.0 | ± 2.4 | 25.8 | ± 0.8 |

The brightness change per voltage increment (|dB/dV_m_|) as a function of V_m_ of VSD-based GEVIs follows a bell-shaped curve and is maximal at V_half_ (Eq. 2, Fig. 1D). For optimal resolution, V_half_ is located in the center of the V_m_ range of interest, which, in the case of mammalian cells, falls between -50 and -25 mV. To shift V_half_ into this target V_m_ range, we reversed a R396Q mutation, which was originally introduced in ASAP1 to shift V_half_ towards negative voltages (20). We found that the Q396R mutation not only shifted V_half_ but also improved B_max_, ΔB_max_ and k_s_ (Fig. 1C and table 1). As a result, the maximum |dB/dV_m_| of ASAP3:Q396R was more than doubled (p = 7.8 × 10^−6^) over ASAP3 (Fig. 1D). However, owing to a too strong right-shift of V_half_ to 10.9 ± 2.1 mV the majority of the overall brightness change occurred outside the physiological resting V_m_ range (Fig. 1C). Interestingly, the down state and the up state of ASAP3:Q396R were brighter compared to ASAP3, which suggests additional VSD configurations. The existence of states below the down and above the up state is supported by a recent study by Shen *et al*. which reports the presence of additional states (up-plus and down-minus) in the VSD of the *Ciona intestinalis* (Ci) VSP (28).

We aimed to maintain the improvements obtained with the Q396R mutation and shift V_half_ into the target V_m_ range. Therefore, we generated a structural model of ASAP3:Q396R that most closely resembled the up state of the VSD (28,40,41) and performed site-saturation mutagenesis based on mKate2-ASAP3:Q396R at three residues (T134, N138, and Y141) in transmembrane segment 3 (Fig. 1E). These residues are in close proximity to two of the voltage-sensing charges in S4 and are therefore likely to influence the voltage-sensor movement (28). Shen *et al*. recently showed that residues T197 and Y200 in the VSD of the Ci-VSP, which correspond to N138 and Y141 in ASAP3, can shift the equilibrium between up and down states, further supporting the choice of these sites for mutagenesis (28). For a systematic performance comparison, we recorded the brightness–voltage relationships of all mutants with sufficient membrane expression (Figs. S1 to S3 and table 1) and determined the maximum |dB/dV_m_| and V_half_ (Fig. 1F). Out of all evaluated mutants we chose N138G for further optimization due to its high maximum |dB/dV_m_| and a V_half_ of -19.1 ± 2.2 mV, which is close to the desired target V_m_ range. We further combined N138G with Y141T, with a V_half_ (-57 ± 3 mV) slightly to the left of the target range. The resulting triple mutant N138G:Y141T:Q396R featured a V_half_ of -42.2 ± 2.0 mV and a maximum |dB/dV_m_| of 0.379 ± 0.010%_EGFP_ mV^-1^ (Fig. 1F).

To further enhance the brightness of the sensor we examined several mutations which have been reported to increase the fluorescence intensity of GFP variants (Fig. S4) (42). Out of the tested mutants only S318A increased the molecular brightness: in the background of N138G:Y141T:Q396R, mutation S318A enhanced B_max_ from 54.3 ± 1.9 to 77.5 ± 1.7%_EGFP_ (p = 8.4 × 10^−6^, Fig. 1F). Due to its ultrasensitive response, particularly in the physiological resting V_m_ range, we named the resulting N138G:Y141T:S318A:Q396R mutant rEstus (Fig. 1F). Compared to ASAP3, the molecular brightness change of rEstus was improved by 3.2-fold (p = 3.3 × 10^−8^) across the resting V_m_ range and 3.3-fold (p = 1.0 × 10^−9^) across the whole physiological V_m_ range (Fig. 1G). At its sensitivity optimum of -32.0 ± 2.4 mV, which is close to the center of the resting V_m_ range (-50 to 0 mV) of non-excitable cells (2), |dB/dV_m_| of rEstus was increased by 4.6-fold over ASAP3 (Fig. 1H).

To evaluate the response kinetics of all generated sensors towards rapid depolarizing stimuli, we combined whole-cell patch clamp with high-speed photometry (Figs. S5 to S7) and applied step-depolarization protocols from -80 to 40 mV for 2 and 512 ms (Fig. 2A) (43). The relative amplitude during a 2-ms depolarizing pulse (F_2 ms_/F_512 ms_) provided a direct measure of the speed. For quantifying the single-peak performance (ΔB_2ms_), we scaled the normalized traces with the absolute molecular brightness change for a voltage step going from -80 to 40 mV (Fig. 2B). We found that the response kinetics of rEstus (37.0 ± 1.3%) towards depolarizing stimuli was about as fast as those of ASAP3 (39.0 ± 1.1%) (p = 0.29, Fig. 2A). Strikingly, the single-peak performance of rEstus (17.0 ± 0.9%_EGFP_) was improved 3.6-fold (p = 1.1 × 10^−5^) over ASAP3 (4.7% ± 0.5%_EGFP_) (Fig. 2B). While some of the other mutants, such as N138Q (F_2 ms_/F_512 ms_ = 47.3 ± 1.0%), exhibited faster response kinetics, rEstus outperformed all other generated GEVIs in terms of single-peak performance (Fig. 2C).

**Fig. 2.**
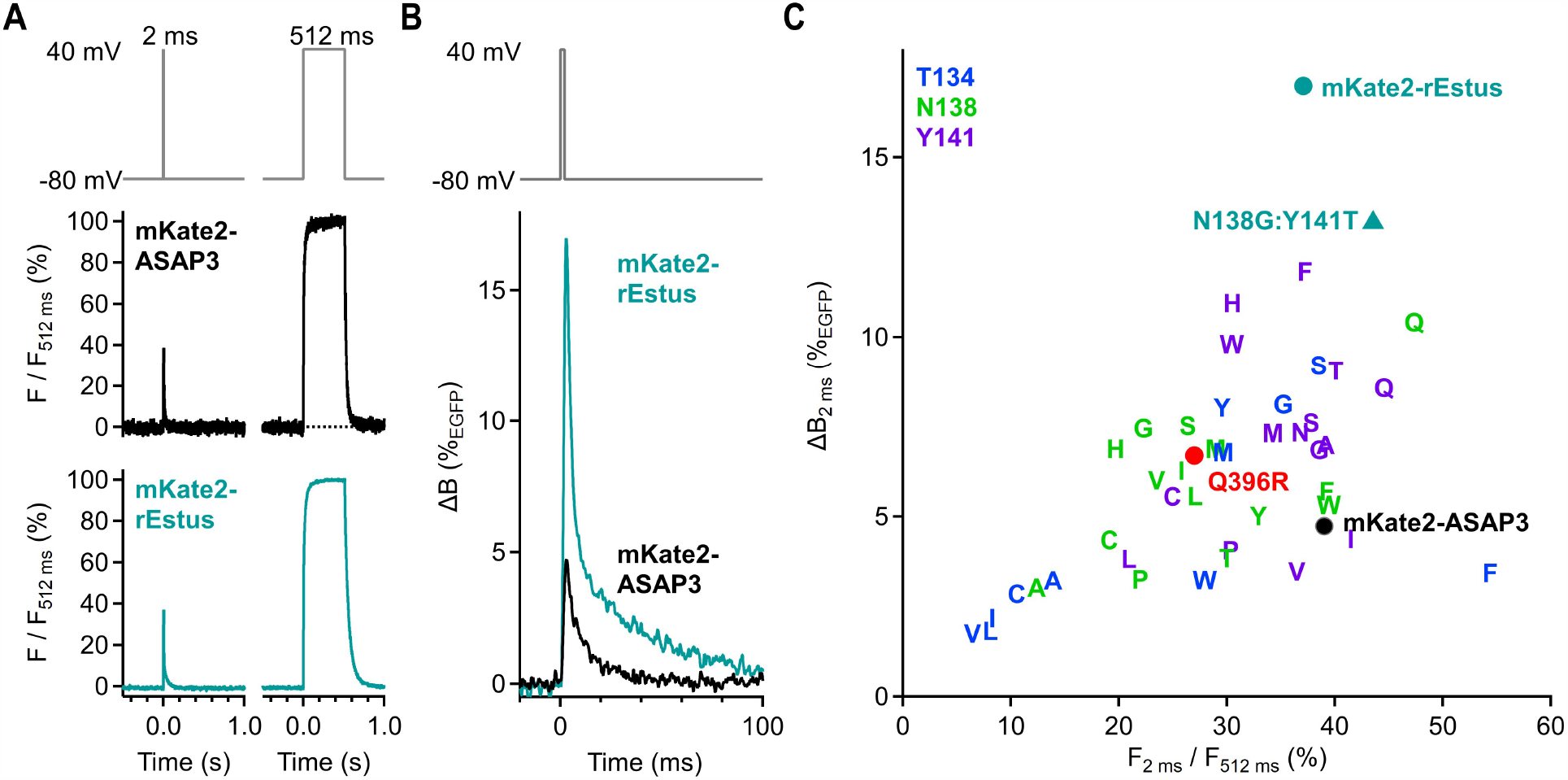
High-fidelity detection of single voltage spikes with rEstus. (**A**) Normalized F_cpGFP_ responses of voltage-clamped HEK293T cells expressing mKate2-ASAP3 or mKate2-rEstus recorded with a photodiode system at 23°C; traces are means with n = 5 each. The relative peak amplitude during the 2-ms pulse (F_2 ms_/F_512 ms_) is a measure of the sensor speed. (**B**) Superposition of the 2-ms responses (from A) scaled by the absolute molecular brightness change at equilibrium. Peak ΔB at 2 ms is a measure of the single-peak performance. (**C**) Single-peak performance plotted against the relative 2-ms peak amplitude for all functional mutants. For n values and average fluorescence traces see Figs. S5 to S7. The analysis of the kinetics for ASAP3 and rEstus across the full voltage range, including depolarizing and hyperpolarizing directions, is shown in Fig. S8.

### Absolute V_m_ measurement with single-protein fluorescence ratiometry

Fusion of mKate2 to a potentiometric GEVI is not only suited to determine the molecular brightness of a sensor molecule but also allows for absolute V_m_ calibration, as we have shown previously (35). However, we observed that the fusion of mKate2 reduced the expression level of the sensor by a factor of 3.1 (Fig. S4D). Some GFP-based sensors do not require a second fluorescence protein for calibration but rather utilize the equilibrium between the neutral (400 nm excitation peak) and anionic (480 nm excitation peak) form of the GFP-chromophore to generate a ratiometric signal (44). We found that rEstus can be calibrated using the same approach (Figs. 3A and B), and, that the ratio of F_cpGFP_ of rEstus excited at 400 and 480 nm (F_480_/F_400_) as a function of V_m_ was remarkably reproducible (Fig. 3C and table 2).

**Fig. 3.**
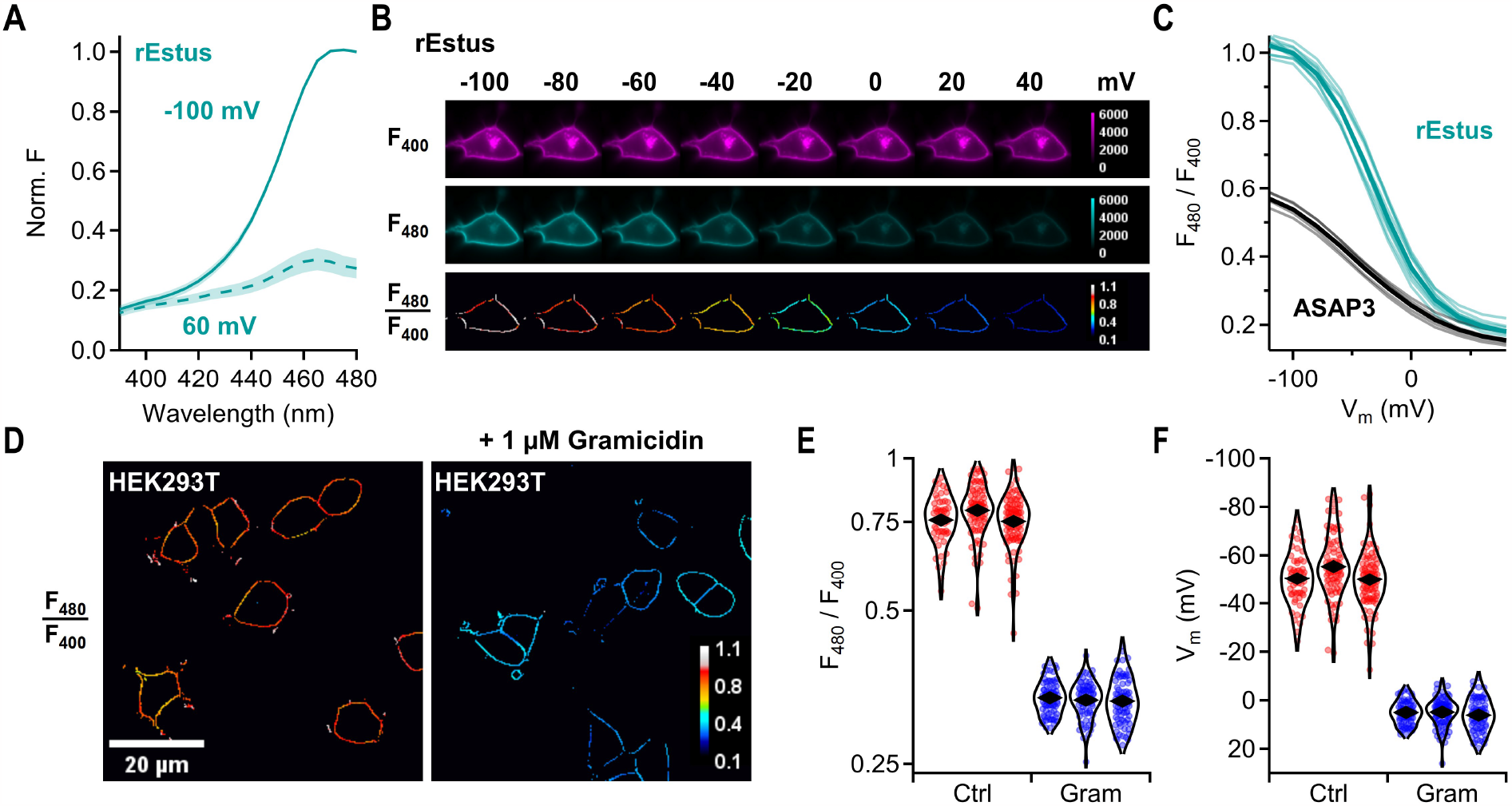
Absolute V_m_ calibration with dual-excitation single-emission ratiometry. (**A**) Normalized excitation spectra of rEstus expressed in HEK293T cells, voltage-clamped to -100 or 60 mV; traces are means of 5 cells, SEM is indicated by shading. (**B**) F_cpGFP_ of a cell expressing rEstus, clamped to the indicated voltages, excited with light of 400 or 480 nm. For F_480_/F_400_ images, only the cell outline was selected using a 3-by-3 ridge filter, to exclude signals from non-responsive intracellular compartments. (**C**) F_480_/F_400_ of rEstus and ASAP3 as a function of V_m_. Thin lines represent recordings from individual cells, thick lines are the mean values; n = 5 for ASAP3 and 10 for rEstus. (**D**) Ridge-filtered F_480_/F_400_ images of HEK293T cells stably transfected with rEstus. Cells on the right were treated with 1 μM of the ionophore gramicidin. (**E**) Violin plots of F_480_/F_400_ of cells with and without 1 μM gramicidin for three cell-culture dishes; the total numbers of analyzed cells were 234 (Ctrl) and 248 (Gram); rhombs represent the means of the distributions. (**F**) Violin plots of V_m_ of the cells from E calculated with the fit parameters (table 2) of the calibration curve in C.

**Table 2.** Parameters of the fits according to Eq. 1 describing the F_480_/F_400_–voltage relationships of rEstus and ASAP3 from Fig. 3.

| GEVI | $n$ | Max.<br>$F_{480}/F_{400}$ | SEM | Max.<br>$\Delta F_{480}/F_{400}$ | SEM | $V_{\text{half}}$<br>(mV) | SEM<br>(mV) | $k_s$<br>(mV) | SEM<br>(mV) |
| --- | --- | --- | --- | --- | --- | --- | --- | --- | --- |
| rEstus | 10 | 1.051 | $\pm 0.008$ | 0.878 | $\pm 0.011$ | -31.9 | $\pm 1.8$ | 25.0 | $\pm 0.3$ |
| ASAP3 | 5 | 0.638 | $\pm 0.010$ | 0.503 | $\pm 0.010$ | -45.7 | $\pm 1.5$ | 39.5 | $\pm 0.8$ |

As a proof of concept, we recorded F_480_/F_400_ from HEK293T cells stably expressing rEstus. We treated cells with 1 μM gramicidin, an ionophore that short-circuits the plasma membrane when integrated (35). The difference in F_480_/F_400_ between control cells and those treated with gramicidin was readily apparent on a single-cell level (Fig. 3D). The average F_480_/F_400_ of untreated cells was 2.3-fold higher (p = 0.7 × 10^−3^, n = 3 culture dishes each) than that of gramicidin-treated cells, indicating depolarization caused by gramicidin (Fig. 3E). Using the parameters from the F_480_/F_400_ calibration curve (table 2), we calculated the corresponding V_m_ for each cell (Fig. 3F): for non-treated HEK293T cells, the average V_m_ was -51.8 ± 1.8 mV, with an inter-cell variability of 11.9 ± 0.6 mV (standard deviation). Gramicidin not only depolarized the cells to 5.4 ± 0.4 mV (p = 0.6 × 10^−3^) but also reduced the inter-cell variability to 5.7 ± 0.6 mV (p = 1.7 × 10^−3^), which indicates that at least part of the inter-cell variability in non-treated HEK293T cells originates from genuine variability in V_m_.

### Endogenous electrical activity of HEK293T cells uncovered with rEstus

The electrical signaling of excitable cells by action potentials with large voltage amplitudes is well established. In contrast, the time course and amplitude of V_m_ variations in non-excitable cells under live-cell conditions are largely unknown (17). To explore the V_m_ dynamics in HEK293T cells we recorded V_m_ using rEstus at 5 Hz (Figs. 4A and B).

**Fig. 4.**
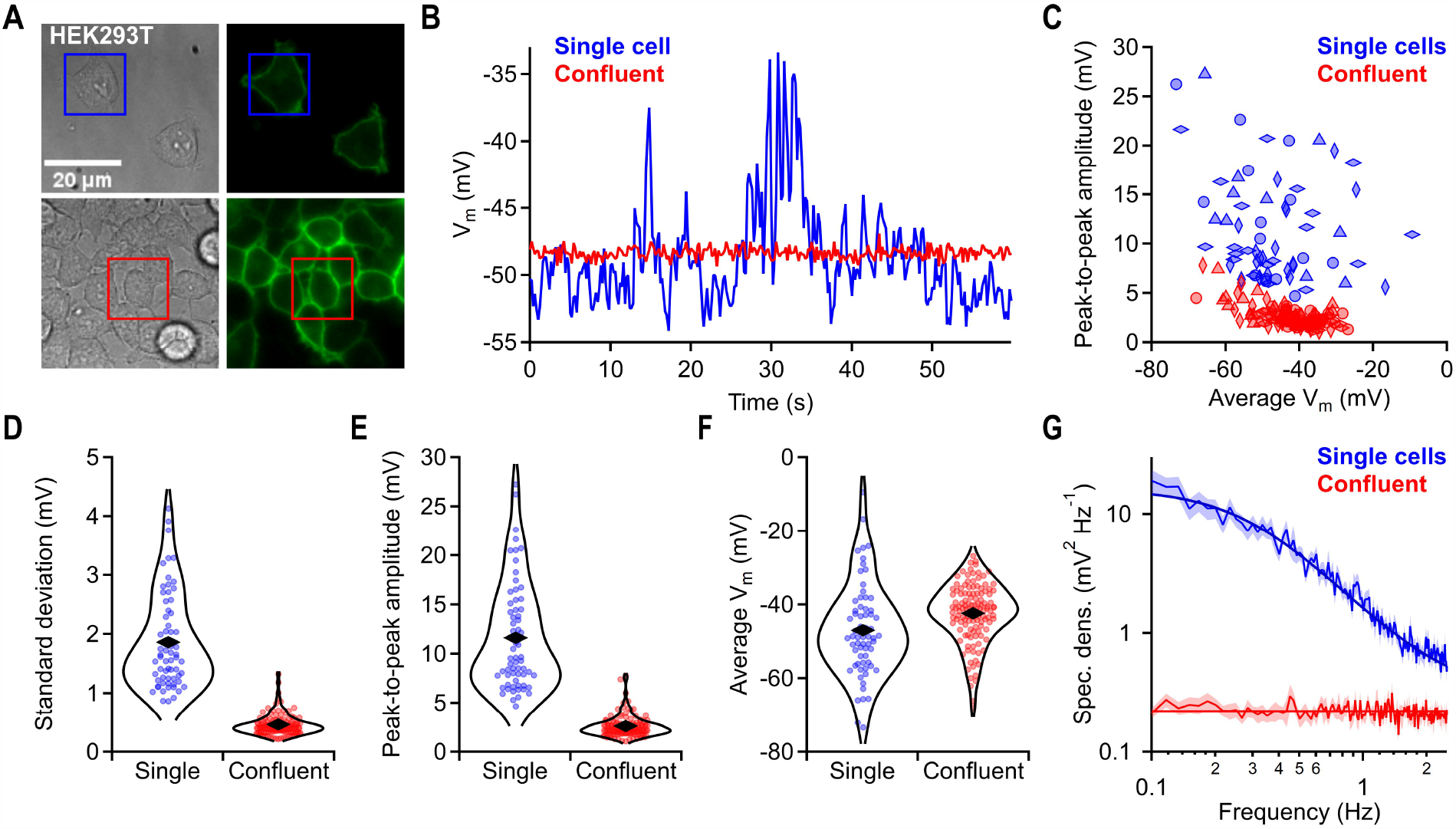
rEstus reveals high V_m_ volatility of individual HEK293T cells. (**A**) Transmission and fluorescence images of HEK293T cells stably expressing rEstus. Cells were either individual or at confluency. V_m_ time-traces of cells with boxes are shown in B. (**B**) Cell-density dependent electrical activity. V_m_ was calculated from F_480_/F_400_ traces of the marked cells in A. (**C**) Peak-to-peak V_m_ amplitudes of 60-s sweeps plotted against the average V_m_ for single cells (n = 65) and cells in confluent clusters (n = 122). Data with same symbol and color are from the same cell culture dish. (**D** to **F**) Violin plots of standard deviation, peak-to-peak amplitude, and average V_m_ (from C). Rhombs are means of the distributions. The data were combined from the four independent cell culture dishes for each condition. (**G**) Mean power spectra of V_m_ data traces from C. Superimposed white noise (0.22 mV^2^/Hz) for confluent cells and white noise (0.3 mV^2^/Hz) plus a square-law filter function (Lorentzian, f_c_ = 1.8 Hz, S = 16 mV^2^/Hz) for individual cells.

Individual HEK293T cells with no connection to other cells featured substantial excursions of V_m_ on timescales spanning from milliseconds to seconds (Fig. 4B). Strikingly, V_m_ fluctuations of cells within a confluent cluster were strongly attenuated. The average size of the V_m_ fluctuations (standard deviation) for individual and confluent cells was 1.89 ± 0.10 mV and 0.46 ± 0.04 mV (p = 0.29 × 10^−3^), the average peak-to-peak amplitudes were 12.0 ± 0.9 mV and 2.57 ± 0.20 mV (p =1.4 × 10^−3^), and the average V_m_ were -47.3 ± 1.0 mV and -42.5 ± 1.8 mV (p = 0.067), respectively (n = 4 culture dishes each, Figs. 4C to F). Thus, while the average V_m_ of single cells and cells within a confluent cluster was similar, V_m_ of individual cells was substantially more volatile. Spectral analysis revealed that V_m_(t) of confluent HEK293T cells predominantly contained white noise across the examined frequency range. In contrast, V_m_(t) of individual cells exhibited greater spectral power, decaying with a square law and a cutoff frequency of approximately 1.8 Hz (Fig. 4G).

### Exploring electrical gap-junction coupling by fluorescence–fluctuation correlation analysis

To infer the electrical coupling of two neighboring cells through gap junctions, it is necessary to measure the electrical resistance between them (45). Since changes in V_m_ are optically resolved using rEstus, the coupling between two cells can be determined by altering the voltage in one cell using whole-cell patch clamp and recording the F_480_ signal in a second cell (cell 1 and 2 in Fig. 5A). Following this approach, we observed that the fluorescence of rEstus in a HEK293T cell connected to the cell in voltage-clamp mode changed by the same amount as in the voltage-clamped cell. This indicated a low cell-to-cell resistance and, hence, a strong electrical coupling between these cells. While electrical coupling between individual cells is detected with the presented method, the throughput is limited by its reliance on electrophysiological techniques. Considering that endogenous electrical fluctuations of HEK293T cells are resolvable with rEstus, it is plausible to investigate electrical connectivity exclusively by analyzing correlated fluorescence fluctuations. In instances of strong electrical coupling, these fluctuations should exhibit a pronounced positive correlation.

**Fig. 5.**
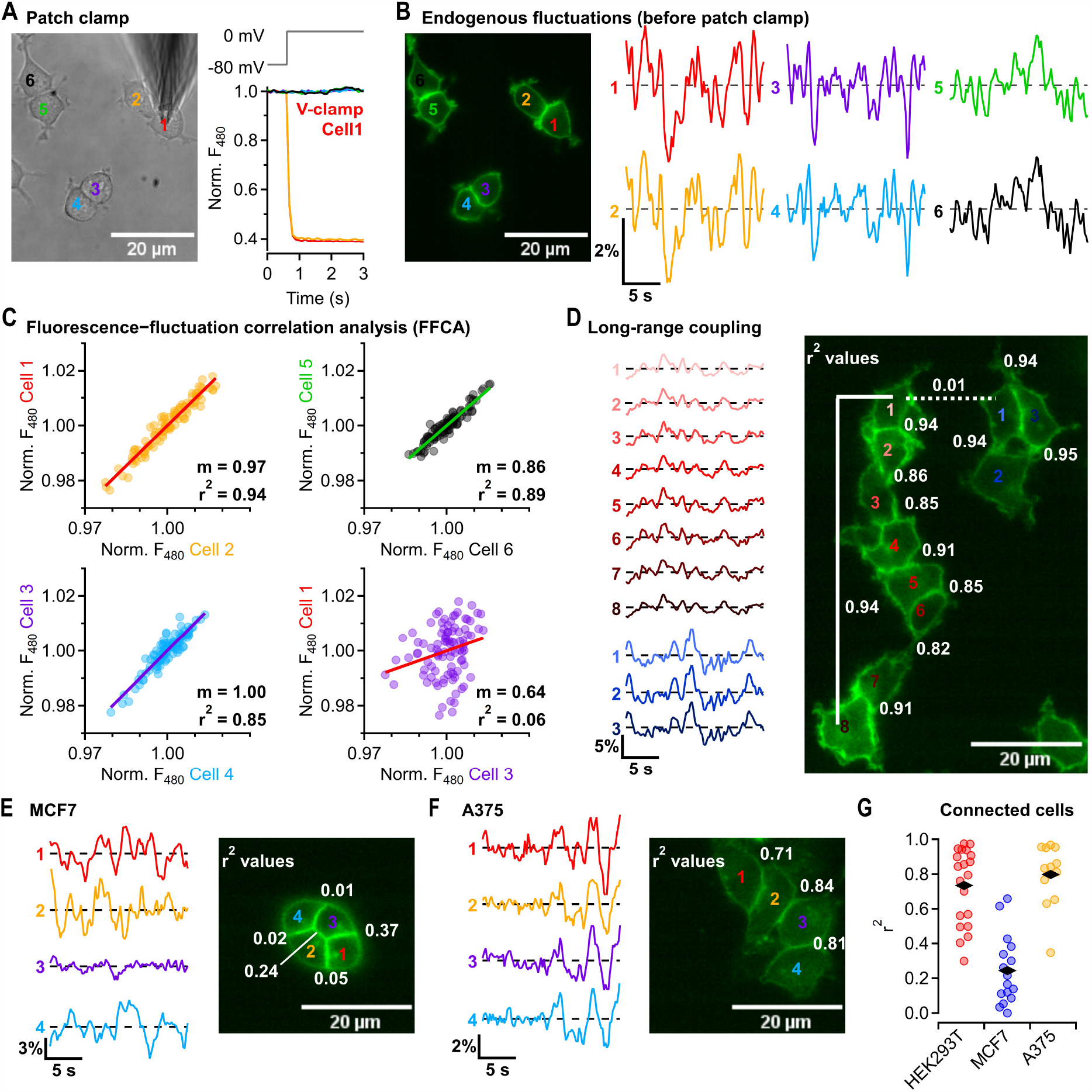
Fluorescence–fluctuation correlation analysis uncovers strong electrical coupling among HEK293T and A375 cells and weak coupling among MCF7 cells. (**A**) (left) Transmission image of HEK293T cells stably expressing rEstus. Cell 1 was in whole-cell patch-clamp configuration. (right) Normalized F_480_ response of all numbered cells to a voltage step. Cells 1 (red) and 2 (orange) were directly connected. (**B**) (left) F_480_ images of HEK293T cell couples shown in (A) before patch clamping. (right) Endogenous F_480_ fluctuations for all six cells. (**C**) Linear correlation of the endogenous F_480_ fluctuations between connected cells (cell 1 and 2, cell 3 and 4, and cell 5 and 6) or non-connected cells (cell 1 and 3). The slope (m) and coefficient of determination (r^2^) for the linear correlation (bold lines) are indicated. (**D**) (left) Endogenous F_480_ fluctuations of two separate clusters of HEK293T cells. The r^2^ values for the corresponding cell-cell connections from fluorescence traces are shown in the fluorescence image (right). The solid line marks the correlation between distant cells within the left cluster, while the dotted line marks the correlation between neighboring cells from separate clusters. (**E**) (left) Endogenous F_480_ fluctuations in human MCF7 breast cancer cells transiently expressing rEstus. The r^2^ values for the corresponding cell-cell connections from fluorescence traces are shown in the fluorescence image (right). (**F**) As in E but for human A375 melanoma cells. (**G**) r^2^ values for neighboring cell pairs. Each data point represents a pair of cells from independent cell clusters with 20 pairs for HEK293T cells, 16 pairs for MCF7 cells, and 13 pairs for A375 cells. Mean values are depicted as rhombs.

Before establishing a whole-cell patch-clamp configuration, we recorded 20-s time traces of F_480_ from the same cells as used for the subsequent patch-clamp analysis (Fig. 5B). All cells within the recorded area exhibited fluctuations in F_480_ with peak-to-peak amplitudes ranging from 2.6% to 4.0%. Remarkably, the F_480_ fluctuations of cells with visible connections (e.g., cell 1 and cell 2) displayed a strong positive correlation with an average coefficient of determination (r^2^) of 0.90 (Fig. 5C). In contrast, for cells without connections (e.g., cell 1 and cell 3), the average r^2^ was only 0.03. We extended the fluorescence– fluctuation correlation analysis (FFCA) to larger clusters (Fig. 5D) and found that the F_480_ fluctuations remained highly positively correlated even between HEK293T cells indirectly connected through several other cells (cell 1 and 8) indicating strong electrical coupling among all cells within the cluster.

To assess whether FFCA is applicable to other cell types, we transiently expressed rEstus in MCF7 breast cancer and A375 melanoma cells. In both cell types, V_m_ was volatile, and the F_480_ signal of rEstus fluctuated on a second timescale, with peak-to-peak amplitudes ranging from 3.5% to 11.7% in MCF7 cells and 4.6% to 6.9% in A375 cells (Figs. 5E and F). In A375 cells, the F_480_ fluctuations in a cluster of cells showed a strong positive correlation (Fig. 5F). In contrast, even in tightly packed clusters of MCF7 cells, individual cells displayed weak correlation with their neighbors or no correlation at all (Fig. 5E). FFCA of additional cell pairs confirmed that both HEK293T (r^2^ = 0.73 ± 0.05) and A375 cells (r^2^ = 0.80 ± 0.05) exhibited similar (p = 0.36) strong electrical coupling whereas MCF7 cells (r^2^ = 0.24 ± 0.05) showed weaker electrical coupling compared to both HEK293T (p = 4.6 × 10^−8^) and A375 cells (p = 1.4 × 10^−8^, Fig. 5G).

## Discussion

In this study, we developed rEstus, a genetically encoded fluorescent voltage indicator with a maximum molecular brightness of 77.5%_EGFP_. When excited at 480 nm, the fluorescence intensity of rEstus changes by around 5-fold across the physiological V_m_ range. rEstus is most sensitive at -32 mV where it achieves a brightness change per voltage increment of 0.55%_EGFP_ mV^-1^. The sensitivity optimum is closely aligned with the average resting V_m_ range (-50 to 0 mV) of proliferating non-excitable mammalian cell types (1,2), such as embryonic and tumor cells, allowing to resolve endogenous sub-second V_m_ variations on the order of ≈1 mV.

The absolute molecular brightness change of rEstus for fast depolarizing voltage spikes is increased 3.6-fold over ASAP3. While the kinetics for depolarizing steps are similar for both sensors, the response to repolarizing voltage steps is slower for rEstus (Fig. S8). This is a disadvantage when resolving high-frequency events, such as trains of action potentials in neurons (34). However, if only the identification of an electrical event is of interest rather than the exact waveform, the slow off-rate of rEstus may be a desirable feature as it prolongs the duration during which the sensor stays in the dim state, thus facilitating the detection of individual action potentials (31,46).

While single-cell V_m_ quantification was achieved with voltage-sensitive dyes using fluorescence-lifetime imaging (47), no GEVI-based approach offered sufficient resolution to quantify the V_m_ of individual cells (35–38). We demonstrated that rEstus can be effectively calibrated using dual-excitation single-emission ratiometry. This calibration method permits single-cell V_m_ measurements with sub-second time resolution. The calibration based on F_480_/F_400_ is also applicable to ASAP3 and likely extends to other ASAP-derived GEVIs.

Probing the coupling of neighboring cells through gap junctions with FFCA using rEstus offers several advantages over methods based on electrodes or transfer of small molecules (48–50). It eliminates the need for additional sample preparation, such as dye loading, and avoids disruption of the cellular environment as in electrode-based methods. In addition, methods that rely on the diffusion of small molecules may not truly reflect the temporal characteristic of the ion flux-mediated electrical connectivity. FFCA provides real-time assessment of the electrical connectivity of large numbers of cells across long distances and facilitates the monitoring of dynamic changes in cell-cell communication. FFCA should be immediately applicable in most laboratories because it only requires a fluorescence microscope equipped with an EGFP filter set and an imaging system supporting a framerate of about 5 Hz. We observed that for large cell populations the endogenous V_m_ fluctuations were attenuated. Hence, for probing electrical connectiveness in large clusters it might become necessary to actively evoke an electrical response, either by standard electrode-based methods or via optogenetic tools, such as channelrhodopsins or membrane targeted micro-/nanoparticles (45,51,52).

We discovered that the V_m_ of HEK293T, A375, and MCF7 cells showed spontaneous fluctuations reminiscent of sub-threshold V_m_ fluctuations in neurons (53). In a recent study, Quicke *et al*. (9) reported similar V_m_ fluctuations in several types of breast cancer using the voltage-sensitive dye di-4-AN(F)EP(F)PTEA. They found that the aggressive MDA-MB-231 cell line was especially electrically active, even in dense layers of cells. However, the sensitivity of di-4-AN(F)EP(F)PTEA is only 6.6% per 100 mV, allowing only for the detection of large voltage excursions. Remarkably, the electrical signals inside these dense layers were only weakly correlated, similar to our observations in MCF7 breast cancer cells.

While the absence of V_m_ fluctuations in HEK293T cells grown to confluency might be explained by changes in the expression/activity patterns of ion channels depending on the cell number (54), V_m_ is stabilized through passive electrical processes. Since every cell membrane acts as an electrical capacitor, the low inter-cell resistance of HEK293T cells connects individual capacitors to a large capacitance. Thus, large groups of electrically coupled cells act as a low-pass filter for high-frequency V_m_ changes. This purely passive electrical process may have functional relevance. A more volatile V_m_ increases the likelihood of spontaneous auto-stimulation events, such as Ca^2+^ influx through voltage-gated Ca^2+^ channels (16,55). Ca^2+^ influx is a stimulus promoting cell proliferation and migration (56,57). Such a negative interrelation between electrical capacitance and Ca^2+^ influx would support auto-stimulation of individual cells and small clusters of cells, which progressively diminishes with an increasing number of cells within a cluster. This may explain how cells can sense the number of other cells within the same cluster, offering a possible mechanism contributing to the well-established contact inhibition of proliferation and contact inhibition of locomotion (58,59). Following this logic, a disturbance of this system, e.g., by the loss of electrical coupling between cells, would cause increased cell migration and proliferation.

While a comprehensive analysis of this working hypothesis is beyond the scope of this study, the provided optical tool rEstus, which permits rapid and quantitative measurements of V_m_ in a non-invasive manner, will help to elucidate the underlying molecular and cellular mechanisms.

## Materials and Methods

### Molecular biology

To derive the molecular brightness from the fluorescence intensity of ASAP3, we generated fusion proteins of mKate2(60) and ASAP3. The construction of the mKate2-ASAP3 plasmid (pCDNA3.1+) involved the linkage of mKate2 (60) to ASAP3 (23) by an *Eco*RI site, flanked by *Nhe*I and *Not*I restriction sites. The mKate2-EGFP construct was created by replacing ASAP3 with EGFP (61). All point mutants based on the ASAP3 template were generated through standard molecular biology methods. The position of each mutation was determined with respect to the N terminus of ASAP3. For mutations in the cpGFP component of the GEVI, the positions are also indicated with respect to the original GFP sequence. Each variant was subsequently validated through sequencing.

For establishing a stable HEK293T cell line, we constructed a pCDNA3.1+/Puro vector by replacing the Kan/Neo cassette in pCDNA3.1+ with the puromycin resistance from pcDNA3.1/Puro-CAG-ASAP1 (52519, Addgene, Watertown, Massachusetts, USA) using *Avr*II and *Bst*BI restriction sites. rEstus-coding DNA was inserted into pCDNA3.1+/Puro using *Nhe*I and *Not*I restriction sites.

### Cell culture

HEK293T cells were obtained from the Centre for Applied Microbiology and Research (CAMR; Porton Down, Salisbury, UK). MCF7 cells were obtained from the European Collection of Authenticated Cell Cultures (ECACC; Porton Down, Salisbury, UK). Both cell lines were cultured in DMEM-F12 (Thermo Fisher Scientific, Waltham, Massachusetts, USA) supplemented with 10% fetal bovine serum (FBS). The cells were maintained in a humidified incubator at 37°C and 5% CO_2_. A375 cells (ATCC, Manassas, VA, USA) were cultured in DMEM (Sigma Aldrich) supplemented with 10% FBS in a humidified incubator at 37°C with 10% CO_2_.

For electrophysiological recordings, 10,000 HEK293T cells were plated per 35-mm glass-bottom dish (Ibidi, Martinsried, Germany) and transfected on the following day with 1 μg plasmid DNA per cell culture dish using the Roti®-fect transfection kit (Carl Roth, Karlsruhe, Germany). MCF7 and A375 cells (ATCC) were transfected with 1 μg DNA using the SF Cell Line 4D-Nucleofector kit (Lonza) following the manufacturer’s instructions via electroporation (4D-Nucleofector, Lonza, Basel, Switzerland). Electrophysiological and imaging experiments were performed one or two days after transfection.

To generate stable HEK293T cell lines, confluent cells from a T75 culture flask were split at a ratio of 1:10. Subsequently, the cells were transfected with a mixture of 10x Roti®-fect and linearized (PvuI) pCDNA3.1+/Puro-rEstus plasmid (10 μg). Three days after transfection, the culture medium was replaced with DMEM-F12 supplemented with 10% FBS and 10 μg/ml puromycin (Gibco). The medium was refreshed every three days for two weeks. The stable cells were further cultivated in the selection medium. Plated cells for fluorescence imaging were kept in medium without puromycin. For experiments with stable cell lines at high and low cell densities, cells were plated at 20,000 or 40,000 cells per 35-mm dish and imaged two or three days after plating.

### Electrophysiological recordings combined with imaging or photometry

Combined electrophysiological and imaging recordings were carried out on an Axio Observer inverted microscope (Carl Zeiss, Jena, Germany) equipped with a Colibri-2 LED illumination system and an EC-Plan Neofluar oil 40x objective (NA 1.3, Zeiss). Images were acquired with a CMOS camera (ORCA-Flash 4.0 Digital Camera C11440, Hamamatsu, Japan) using SmartLux software (HEKA Elektronik, Lambrecht, Germany). Electrophysiological recordings were obtained with an EPC10 double patch-clamp amplifier controlled by PatchMaster software (both HEKA Elektronik). Visualization of mKate2 fusion proteins was achieved with an EGFP/mCherry (59022 Chroma) dual-emission filter set. Excitation of cpGFP was achieved with a 470-nm LED (BP 460/50) with a measured power of 0.709 mW, while excitation of mKate2 was performed with a 530-nm LED (BP 545/40) with a measured power of 1.071 mW.

F_480_/F_400_ was recorded using an EC-Plan Neofluar oil-immersion 40x objective (1.3 NA, Zeiss), an LPXR 495 beamsplitter (Chroma), and a BP 525/50 (Delta Optical Thin Film) emission filter. Excitation was performed using a 405-nm LED (Zeiss) with a BP 400/10 filter (Thorlabs) with a measured power of 0.424 mW, and a 470-nm LED (Zeiss) with a BP 480/10 filter (Thorlabs) with a measured power of 0.126 mW. The light power was measured above the objective.

For measurements of sensor kinetics, whole-cell patch clamp was combined with photometry. Excitation light came from a 470-nm LED (M470L1, Thorlabs), passed through an EC-Plan Neofluar oil-immersion 40x objective (1.3 NA, Zeiss). The filter cube contained a modified filter set 38 (Zeiss) with the excitation filter replaced by a 492/SP filter (Semrock, Rochester, NY, USA). The emitted light was detected with a photodiode (FDU photodiode with viewfinder, T.I.L.L. Photonics, Gräfelfing, Germany) and recorded using PatchMaster software (HEKA Elektronik) at 20 kHz.

For excitation spectra, HEK293T cells expressing variants of ASAP3 were prepared following the previously described protocol. Cells were held under voltage-clamp conditions in the whole-cell configuration. The cells were placed beneath a 40x EC Plan Neofluar oil objective (1.3 NA, Zeiss), and images were acquired using a ProgRes MF cool camera (JenOptik, Jena, Germany) in conjunction with the modified filter set 38 (Zeiss) mentioned above. An XBO lamp coupled to a monochromator (Polychrome V; T.I.L.L. Photonics) served as the excitation source. Images were acquired for each wavelength with a fixed exposure time of 200 ms. The excitation spectra were corrected based on the light intensity at each wavelength. Furthermore, the spectra were normalized to the fluorescence intensity of an image of mKate2 obtained using filter set 14 (Zeiss) and excitation at 550 nm, following the configuration mentioned above.

Patch-clamp pipettes were fabricated from borosilicate glass, coated with dental wax, and fire-polished to obtain resistances of 1-2 MΩ. Before recording, the culture medium was replaced with an external solution containing 146 mM NaCl, 4 mM KCl, 2 mM CaCl_2_, 2 mM MgCl_2_, and 10 mM HEPES; pH 7.4 at 23°C was adjusted with NaOH. The pipette solution contained 130 mM KCl, 2.5 mM MgCl_2_, 10 mM EGTA, and 10 mM HEPES; pH 7.4 at 23°C was adjusted with KOH.

### Live-cell fluorescence imaging

For live-cell experiments, transfected cells were washed once with 1 ml external solution supplemented with 5 mM glucose and subsequently maintained in 2 ml of the same solution. For experiments with gramicidin, 1 μl of a 2 mM stock solution of Gramicidin D (G5002, Sigma-Aldrich) in ethanol was added to 2 ml of external solution with 5 mM glucose. Imaging of the cells was conducted 15 min after the medium exchange and continued for a maximum duration of 15 min at 23°C.

F_480_/F_400_ images were acquired with an Axio Observer inverted microscope equipped with a Colibri-2 LED illumination system and an EC-Plan Neofluar oil 40x objective (NA 1.3, Zeiss). For slow imaging experiments (Fig. 3), six F_480_/F_400_ images were acquired with a 1-s exposure for each wavelength. The first image was discarded to allow F_480_/F_400_ to stabilize, and the average F_480_/F_400_ of the remaining five images was used for analysis. For high-speed F_480_/F_400_ imaging experiments (Figs. 4 and 5) images were acquired either at 20 Hz (with 35 ms exposure time for each wavelength) and filtered at 10 Hz or recorded at 10 Hz (with 90 ms exposure time for each wavelength). The excitation light came from 405-nm (0.424 mW) and 470-nm LEDs (0.126 mW) with BP 400/10 and BP 480/10 excitation filters, respectively; the filter cube further contained an LPXR 495 beamsplitter and a BP 525/50 emission filter.

### 3D model of ASAP3:Q396R

The 3D model of ASAP3:Q396R was generated *in silico* using ColabFold (62). We aligned the model with the crystal structure VSD of the Ci-VSP in the up and down state and found that it most closely resembles the up state of the VSD (not shown) (41). As the prediction did not account for the autocatalytic generation of the chromophore, the model was aligned with the crystal structure of gCaMP2 using UCSF Chimera (63). In all depictions the inner helix of the beta-barrel structure of the cpGFP from gCaMP2 is shown (64).

### Data analysis

All raw image sets were analyzed using Fiji software (65). For the analysis of widefield fluorescence images, regions of interest (ROIs) were delineated along the cell periphery, excluding areas with poor plasma membrane localization. The extracted data were then background-corrected for each fluorescence channel. The quality of V_m_ calibration strongly depended on the ROI selection. We observed that F_480_ exhibited better apparent targeting to the plasma membrane than F_400_. The F_400_ signal generated, in addition to the plasma membrane signal, bright spots inside the cell, presumably from endosomes and Golgi, which are dimmer or absent in the F_480_ signal.

To highlight the membrane-specific signal in the pixel-by-pixel ratio images (Fig. 3), we applied four 3-by-3 convolution filters to the F_480_ image. For each pixel, the maximum value among the four filtered images was selected to construct a ridge-filtered image. Subsequently, an Otsu thresholding procedure was applied to the filtered image to generate a binary mask (65). This mask was then applied to the corresponding pixel-by-pixel ratio image, effectively isolating the outline of the cells.

Convolution filters:

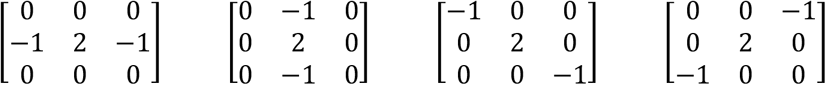

To determine the relative molecular brightness (B) of the cpGFP signal compared to EGFP, the F_mKate2_/F_cpGFP_ values of the mKate2 fusion proteins were normalized to the average F_mKate2_/F_EGFP_ value (4.01 ± 0.05, n = 168 cells) obtained from an mKate2-EGFP fusion protein with an equivalent exposure time ratio. This normalization allowed for a quantitative assessment of the molecular brightness of cpGFP relative to EGFP.

The relationships of molecular brightness versus membrane voltage (V_m_) were fitted using a Boltzmann-type equation:

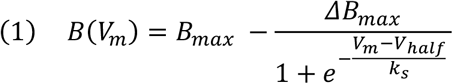

The parameters were as follows: the maximum molecular brightness (B_max_), the maximal change in molecular brightness (ΔB_max_), the voltage at which half-maximal brightness change occurred (V_half_), and the steepness of the response curve (k_s_). The fit parameters for all constructs are given in table S1. For F_480_/F_400_(V), B_max_ and ΔB_max_ in Eq. 1 are replaced by maximal F_480_/F_400_ and maximal change in F_480_/F_400_, respectively.

The brightness change per voltage increment is described by the first derivative of Eq. 1:

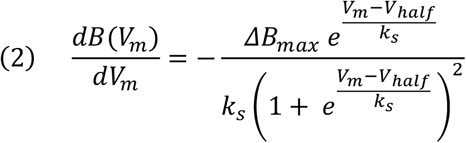

The brightness change per voltage increment of each sensor was calculated for V_m_ = V_half_ in Eq. 2.

For characterization of the sensor kinetics, the absolute changes in molecular brightness for a 2-ms pulse from -80 to 40 mV were calculated by multiplying the equilibrium brightness change between 40 mV (B(40 mV)) and -80 mV (B(-80 mV)) with the ratio of the peak fluorescence during the 2-ms pulse (ΔF_2 ms_) and the average F_cpGFP_ between 430 and 480 ms of the 512-ms pulse:

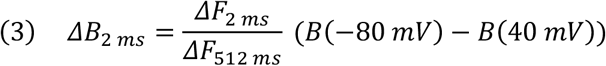

The kinetics of the normalized fluorescence changes in response to voltage steps were analyzed with double-exponential fits:

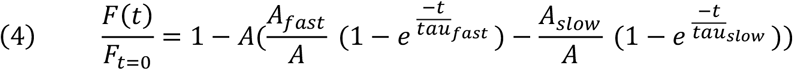

For quantitative high-speed recordings, the F_480_/F_400_ signal was converted into a voltage using the following calibration curve parameters: maximal F_480_/F_400_ 1.051 ± 0.008, maximal change in F_480_/F_400_ 0.878 ± 0.011, V_half_ -31.9 ± 1.8 mV, and k_s_ 25.0 ± 0.3 mV.

For the correlation analysis of neighboring cells, only normalized F_480_ signals were used. ROIs were delineated across the cell body, ensuring no overlap of membrane areas between neighboring cells. The fluctuations of neighboring cells with visible connections were analyzed by linear correlation of a 20-s time trace recorded at 10 Hz and resampled at 5 Hz. Because of the observed photoswitching behavior of ASAP derivatives (66), the first second was not considered for analysis to allow the F_480_/F_400_ signal to stabilize. The traces were linearly corrected for photobleaching. Electrical connectivity between neighboring cells was quantified by calculating the Pearson’s coefficient of determination (r^2^).

### Statistical Analysis

Data are presented as means ± SEM, unless specified otherwise. The number of individual measurements is denoted as n. For statistical comparisons of samples, two-tailed Student’s t-test were applied; resulting p values are provided in the text.

## Acknowledgments

We thank Angela Roßner, Silke Tonndorf-Martini, and Prof. Dr. Markus Gräler for technical support.

## Funding

Simons Foundation Autism Research Initiative (SFARI) (SHH, 705944SH)

German Academic Exchange Service (AGN, 91819480).

## Author contributions

Conceptualization: PR, RS, SHH

Methodology: PR

Investigation: PR, AGN, NG, SNCW, LM

Formal Analysis: PR

Visualization: PR

Supervision: PR, RS, SHH

Writing—original draft: PR

Writing—review & editing: PR, RS, SHH

Funding Acquisition: SHH

## Data and materials availability

All data are available in the main text or the supplementary materials. The authors agree to make the data and materials supporting the results or analyses presented in the paper available upon reasonable request. The rEstus expression vector is available from Addgene (209767).

