## Supplemental information for "An ultrasensitive genetically encoded voltage indicator uncovers the electrical activity of non-excitable cells"

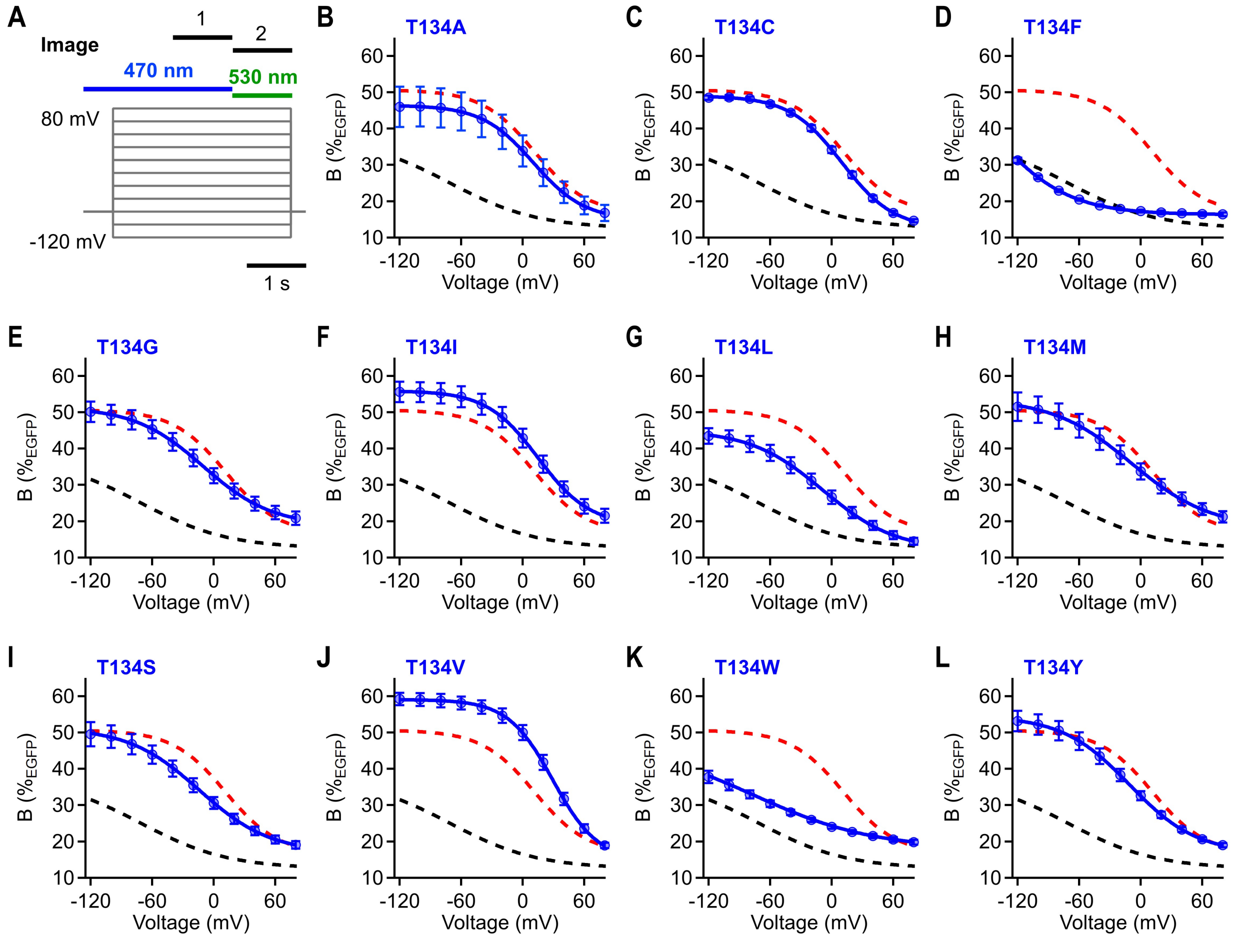


**Fig. S1. Molecular brightness–voltage relationship of ASAP3:Q396R:T134 variants**

(**A**) Imaging protocol for all molecular brightness–voltage recordings. Light of 470 nm and 530 nm wavelength was applied as indicated. Excitation with 470 nm started 1.5 seconds before image acquisition to measure fluorescence after the internal photoswitching of cpGFP. Images for cpGFP and EGFP were acquired with 470 nm illumination, and for mKate2, 530 nm illumination was used. The acquisition was performed using an EGFP-mCherry filter set (Chroma). (**B** to **L**) Molecular brightness–voltage relationships of all mKate2-ASAP3:Q396R:T134 mutants with sufficient plasma membrane expression. The molecular brightness values were obtained by normalizing the recorded $F_{cpGFP}/F_{mKate2}$ signals to the average $F_{EGFP}/F_{mKate2}$ of mKate2-EGFP (4.01 ± 0.05, n = 168 cells) expressed in HEK293T cells and recorded under the same illumination conditions. The mean fits of the mKate2-ASAP3 (black) and mKate2-ASAP3:Q396R (red) brightness–voltage relationships are shown as dashed lines. The data are presented as means with standard error of the mean (SEM) and superimposed fits according to Eq. 1. For n and fit parameters, see table 1.

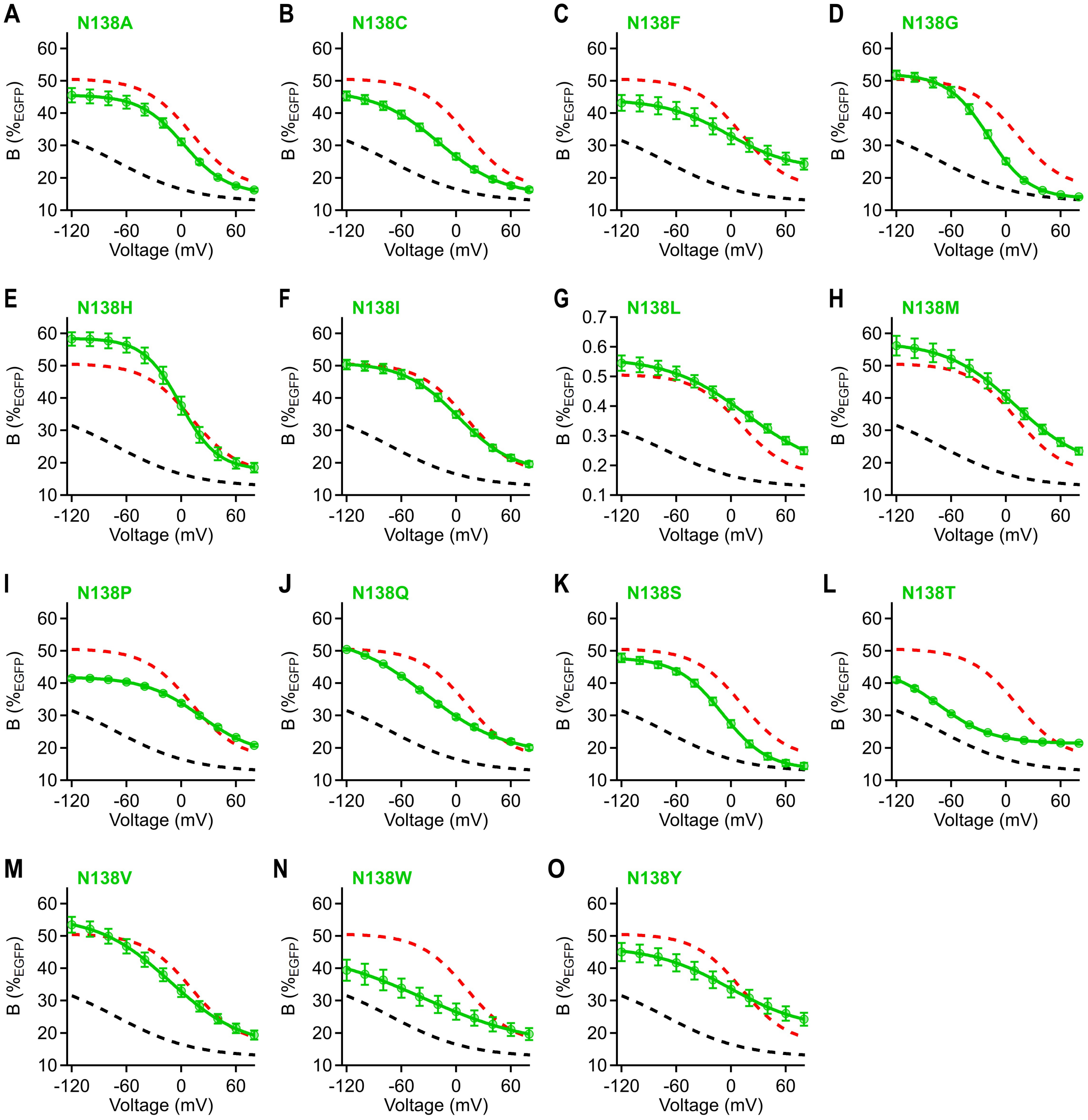


**Fig. S2. Molecular brightness–voltage relationship of ASAP3:Q396R:N138 variants**

(**A** to **O**) Molecular brightness–voltage relationships of all mKate2-ASAP3:Q396R:N138 mutants with sufficient plasma membrane expression. Data were acquired and analyzed as described in Fig. S1. For n and fit parameters, see table 1. The mean fits of the mKate2-ASAP3 (black) and mKate2-ASAP3:Q396R (red) brightness–voltage relationships are shown as dashed lines.

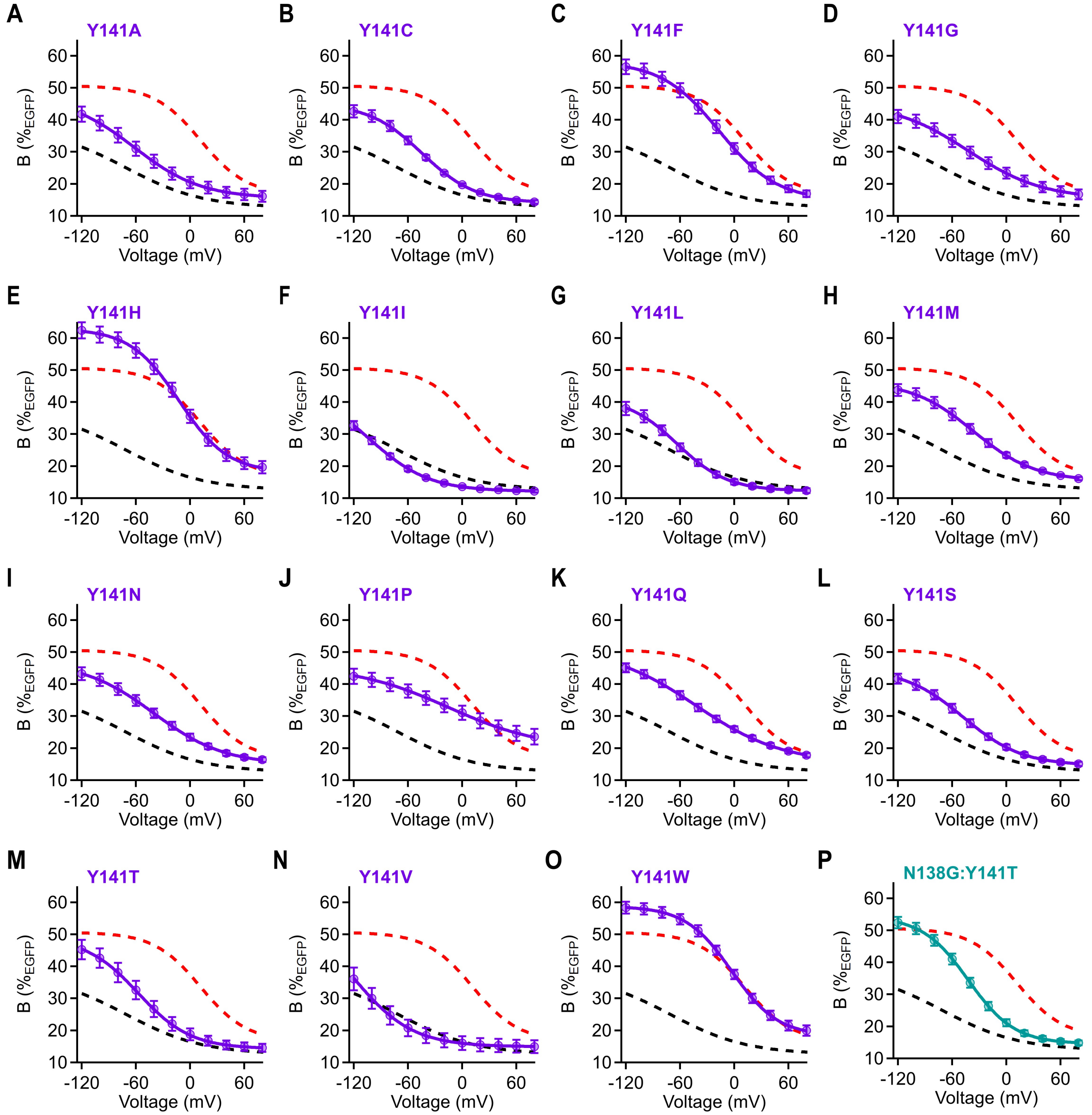


**Fig. S3. Molecular brightness–voltage relationship of ASAP3:Q396R:Y141 variants**

(**A** to **P**) Molecular brightness–voltage relationships of all mKate2-ASAP3:Q396R:Y141 mutants with sufficient plasma membrane expression. Data were acquired and analyzed as described in Supplementary Fig. 1. For n and fit parameters, see table 1. The mean fits of the mKate2-ASAP3 (black) and mKate2-ASAP3:Q396R (red) brightness–voltage relationships are shown as dashed lines.

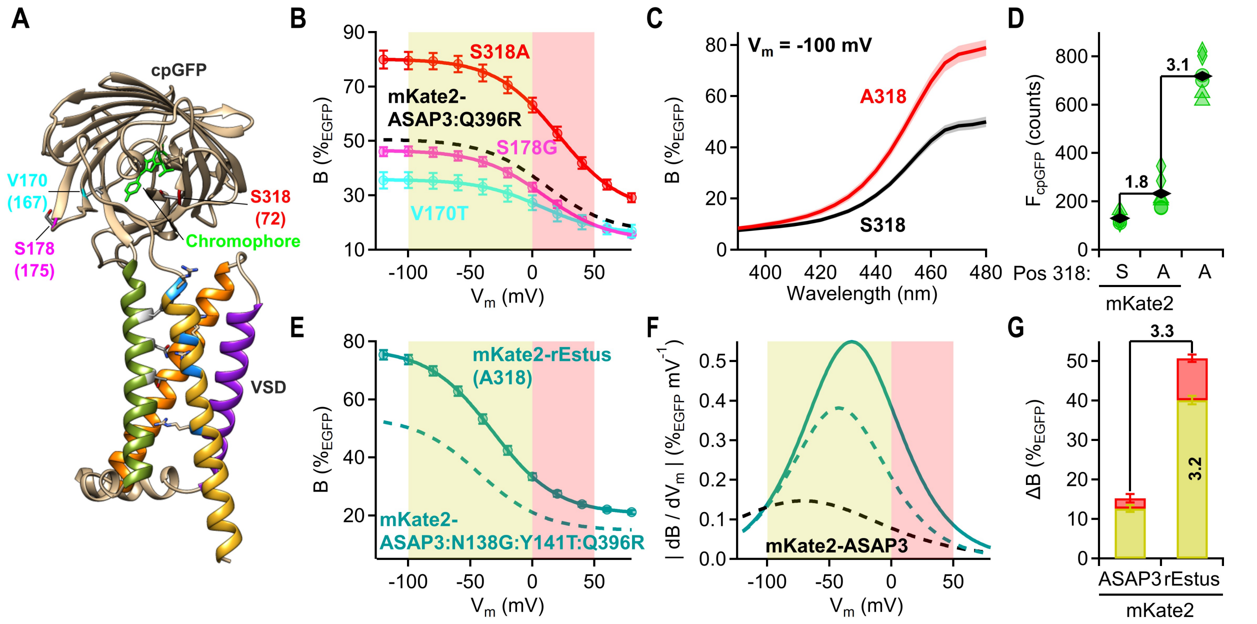


**Fig S4. Mutation S318A in cpGFP increases the molecular brightness of ASAP3 derivatives by up to 60%**

(**A**) 3D model of ASAP3:Q396R. Mutated residues in the cpGFP moiety are numbered with respect to ASAP3 and, in parentheses, with respect to GFP. (**B**) ($V_m$) relationship of mKate2-ASAP3:Q396R variants S318A (red), S178G (magenta), and V170T (cyan), recorded from HEK293T cells. The fit curve of mKate2-ASAP3:Q396R from Fig. 1C is shown for comparison (dashed line). (**C**) Excitation spectra of cpGFP of mKate2-ASAP3:Q396R (S318, n = 5) and the S318A variant (S318A, n = 4) in HEK293T cells voltage-clamped at -100 mV; data are mean ± SEM. The spectra were normalized to the mKate2 signal and scaled to reach the same maximum B($\%_{EGFP}$) for S318 as in B. (**D**) Absolute green fluorescence intensity ($F_{cpGFP}$) of HEK293T cells expressing ASAP3:Q396R and the S318A variant with and without N-terminally fused mKate2. Individual green symbols represent medians of 3700-7400 cells, and each type of symbol represents measurements from one day. Black rhombs represent mean values of the individual medians. (**E**) B($V_m$) relationship of mutant mKate2-ASAP3:N138G:Y141T:S318A:Q396R, termed mKate2-rEstus. For reference, B($V_m$) for the construct with S318 is shown as a dashed line. (**F**) Absolute value of the first derivative (from E) as a function of $V_m$. For reference, $|dB/dV_m|$ is shown for mKate2-ASAP3:N138G:Y141T:Q396R and mKate2-ASAP3 (black) as dashed lines. (**G**) Absolute change in molecular brightness (ΔB) across the resting $V_m$ (yellow) and overshoot (red) range for mKate2-ASAP3 and mKate2-rEstus. Numbers indicate the gain in molecular brightness change in the indicated ranges of mKate2-rEstus with respect to mKate2-ASAP3. All data were obtained from voltage-clamped HEK293T cells and are presented as means ± SEM. For more details, such as n values and fit parameters, refer to table 1.

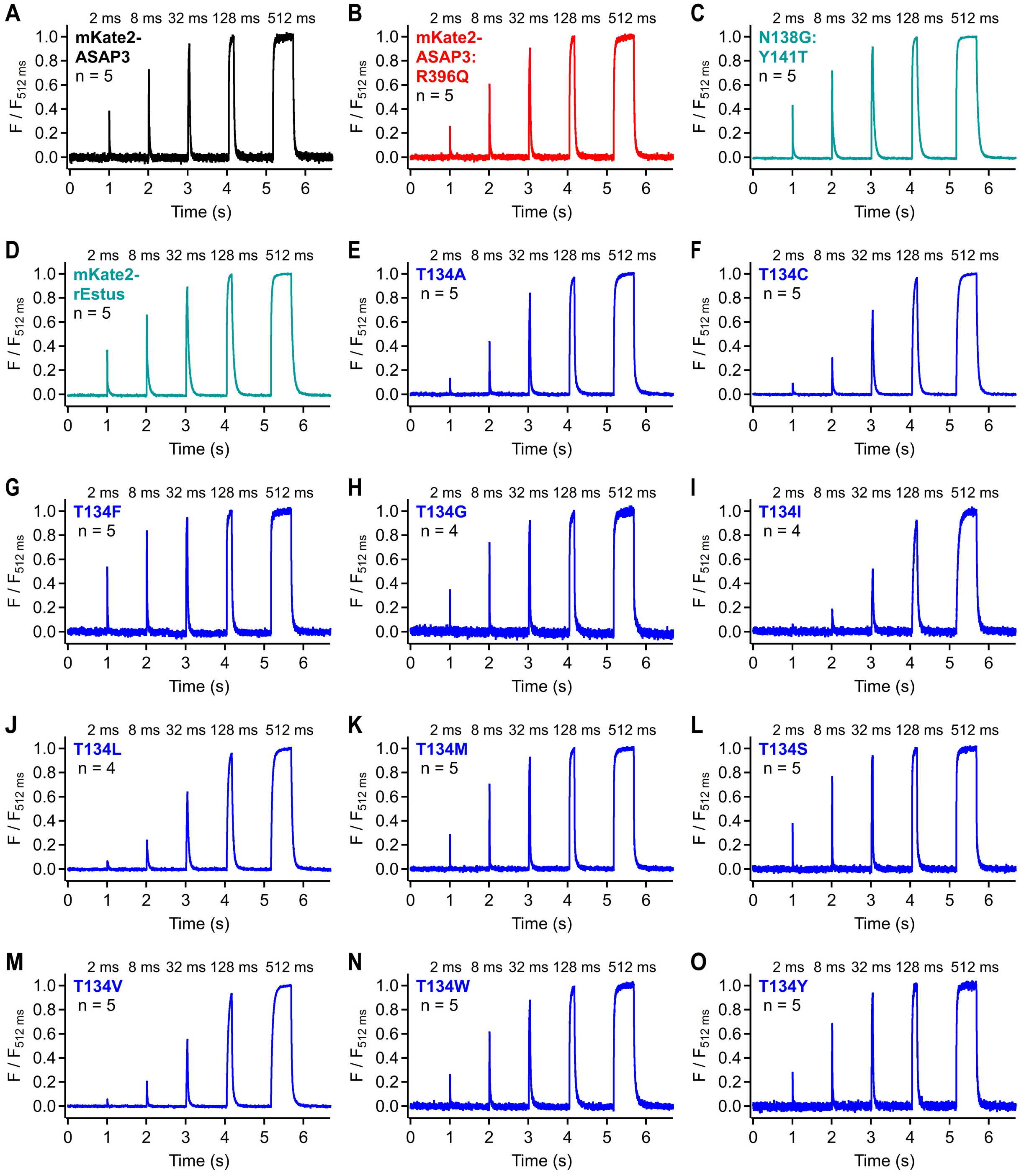


**Fig S5. Voltage-sensor kinetics**

**(A** and **B)** $F_{cpGFP}$ responses of voltage-clamped HEK293T cells expressing mKate2-ASAP3 or mKate2-ASAP3:Q396R recorded with a photodiode system. Voltage steps of the same amplitude (-80 mV holding potential to 40 mV), but with varying durations (ranging from 2 ms to 512 ms), were applied to cells expressing the specified constructs. The traces are the means of the indicated number of recordings from independent cells. The traces were linearly corrected for bleaching, and normalized to the average $F_{cpGFP}$ signal between 430 and 480 ms of the 512-ms pulse. (**C** to **O)**, As in A and B but for cells expressing derivatives of mKate2-ASAP3:Q396R.

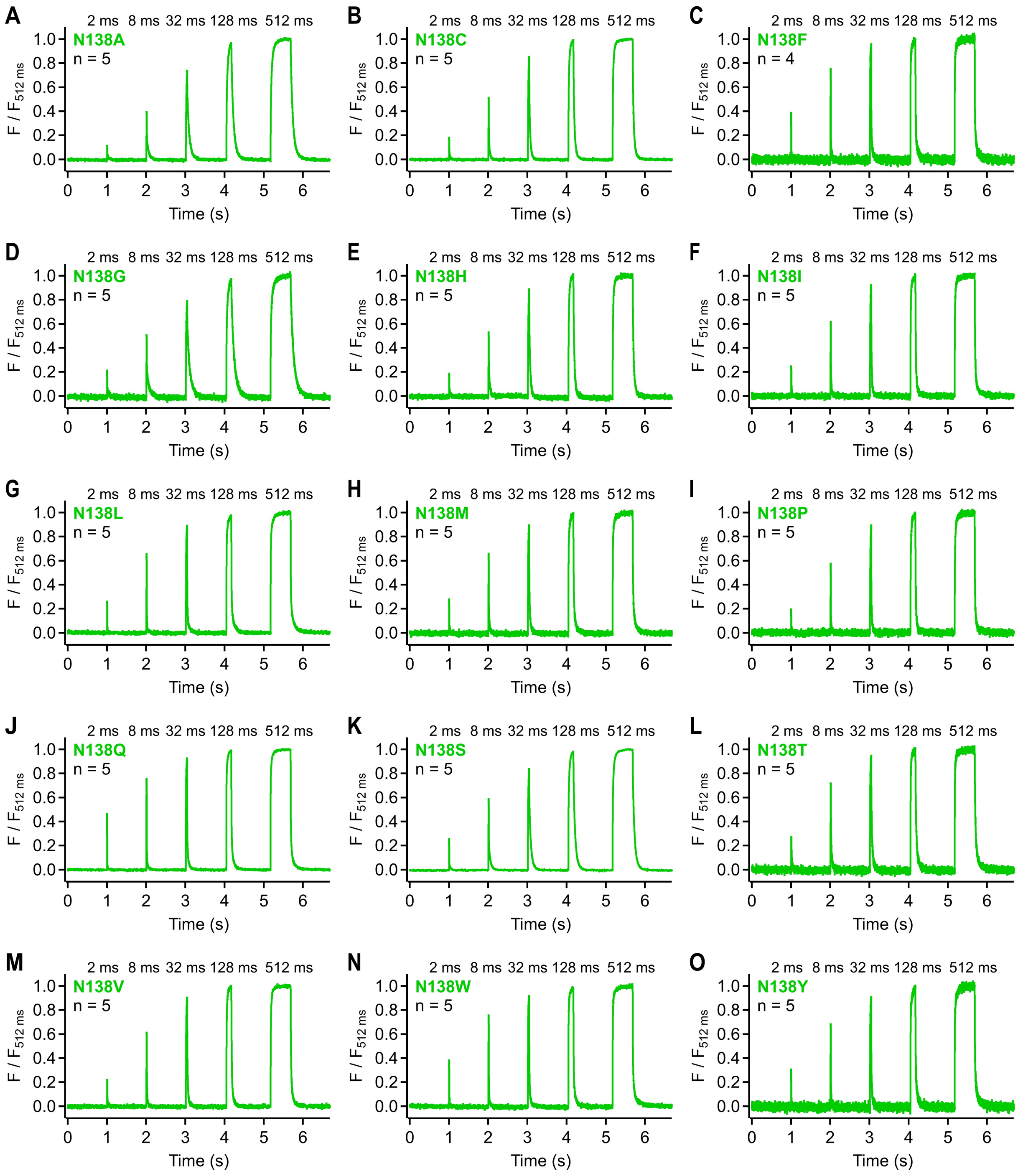


**Fig. S6. Voltage-sensor kinetics for ASAP3-Q396R:N138 variants**

(**A** to **O**) $F_{cpGFP}$ responses of voltage-clamped HEK293T cells expressing N138 derivatives of mKate2-ASAP3:Q396R recorded with a photodiode system. Voltage steps of the same amplitude (-80 mV holding potential to 40 mV), but with varying durations (ranging from 2 ms to 512 ms), were applied to cells expressing the specified constructs. The traces are the means of the indicated number of recordings from independent cells. The traces were linearly corrected for bleaching, and normalized to the average $F_{cpGFP}$ signal between 430 and 480 ms of the 512-ms pulse.

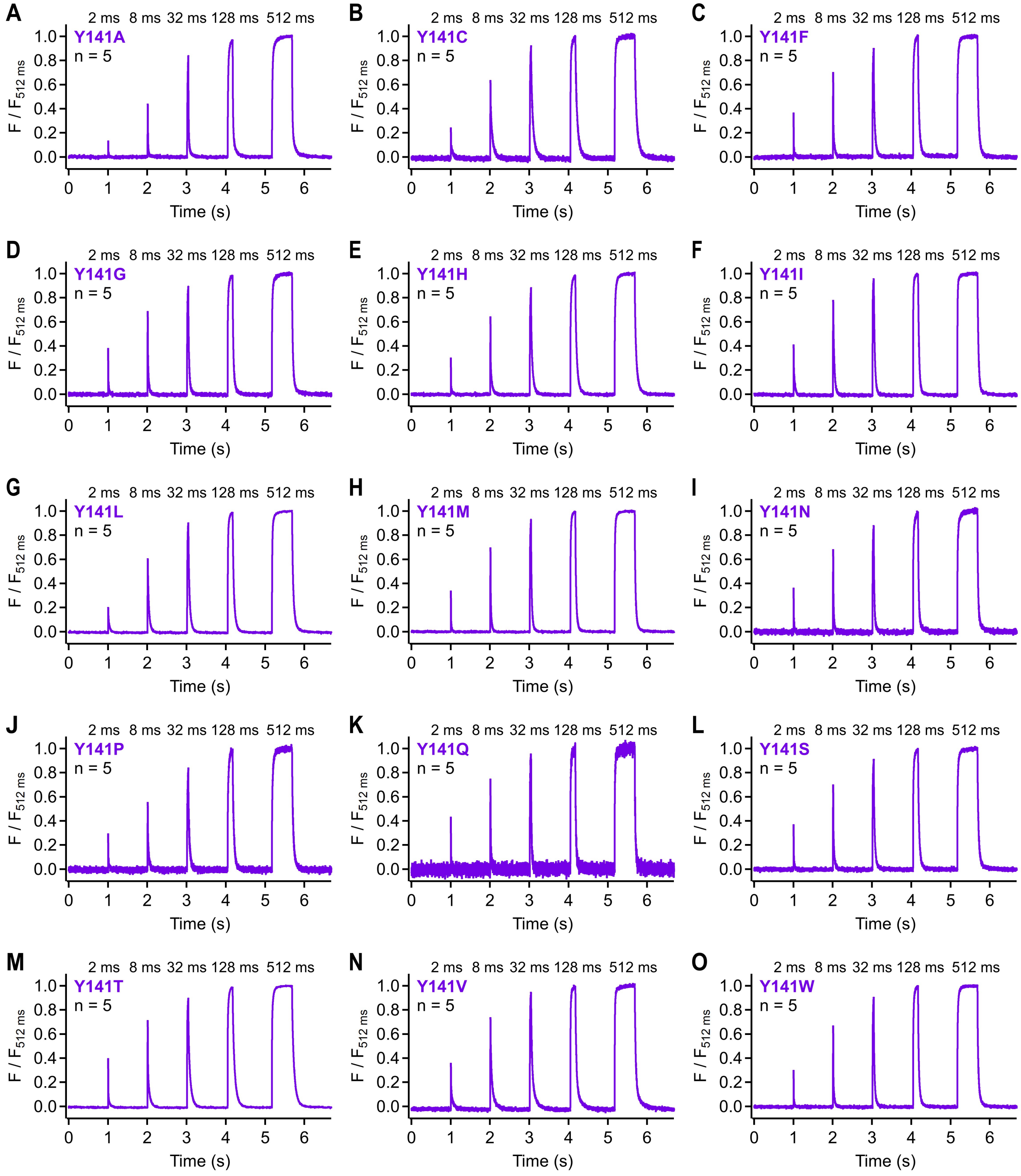


**Fig. S7. Voltage-sensor kinetics for ASAP3-Q396R:Y141 variants**

(**A** to **O**) $F_{cpGFP}$ responses of voltage-clamped HEK293T cells expressing Y141 derivatives of mKate2-ASAP3:Q396R recorded with a photodiode system. Voltage steps of the same amplitude (-80 mV holding potential to 40 mV), but with varying durations (ranging from 2 ms to 512 ms), were applied to cells expressing the specified constructs. The traces are the means of the indicated number of recordings from independent cells. The traces were linearly corrected bleaching, and normalized to the average $F_{cpGFP}$ signal between 430 and 480 ms of the 512-ms pulse.

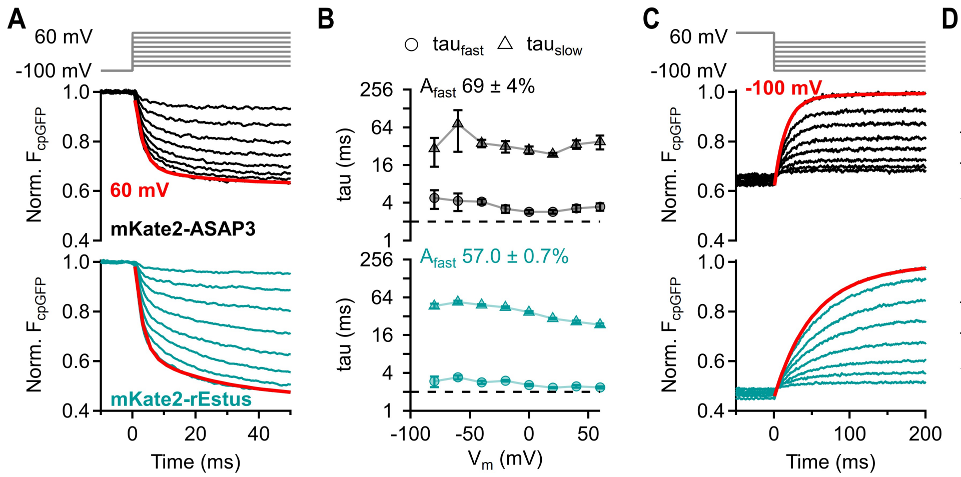


**Fig. S8. Voltage dependence of rEstus and ASAP3 kinetics**

(**A**) Normalized $F_{cpGFP}$ responses of voltage-clamped HEK293T cells expressing mKate2-ASAP3 or mKate2-rEstus to the indicated depolarizing voltage steps recorded with a photodiode system at 23°C; all traces were fit with a double-exponential function (Eq. 3, shown in red for a step to 60 mV). (**B**) The fast and slow time constants from the time-course fits (from A) plotted as a function of $V_m$. The means ± SEM of the relative amplitudes of the fast component ($A_{fast}$) are also indicated; straight lines connect points for clarity; n = 4 for mKate2-ASAP3 and n = 5 for mKate2-rEstus. (**C** and **D**) As in (A and B) but for repolarizing voltage steps; fit curves in C are shown for a step to -100 mV.

### Supplementary Methods

#### Live-cell fluorescence imaging

For expression analysis (Fig. S4D), HEK293T cells were seeded at a density of 20,000 cells per 35-mm dish and transfected with a mixture of 0.75 µg of mKate2-ASAP3:Q396R, mKate2-ASAP3:S318A:Q396R, or ASAP3:S318A:Q396R (all in pCDNA3.1(+)) plasmids along with 0.25 µg of pCNDA3.1-mCherry using ROTI®Fect transfection reagent. The cells were imaged two days after transfection.

Imaging experiments were conducted using an Eclipse-Ti fluorescence microscope equipped with a DS-Qi2 camera (14-bit; Nikon, Tokyo, Japan) and an X-Cite 120 LED light source (Excelitas Technologies, Waltham, Massachusetts, USA), both controlled with NIS4.6 software (Nikon). The temperature of the incubation chamber (Oko Lab, Pozzuoli, Italy) was 37°C. Prior to the measurements, the cell-culture medium was aspirated, and the cells were washed once with 1 ml of external solution supplemented with 5 mM glucose. Subsequently, the medium was replaced with 2 ml of measurement buffer supplemented with 5 mM glucose and 10 µg/ml Hoechst 33342 (Invitrogen). The cells were then incubated for 30 min at 37°C and 5% $CO_2$ before image acquisition. For each 35-mm glass-bottom dish, images were captured from 16 distinct spots using a 10x objective (Plan Apo λ, NA 0.45, Nikon) and an automated stage (H117 stage with a Proscan III controller; Prior Scientific, Cambridge, UK) controlled by NIS 4.6 software (Nikon). The z-focus was maintained stable using the Nikon Perfect Focusing System. The cpGFP signal was obtained using a modified EGFP filter set (GFP-3035D-000), wherein the excitation filter was replaced with a BP 480/10 filter (Thorlabs); the image exposure time was 1 s. The mCherry signal was acquired using a 4040C-000 BrightLine filter with 50 ms exposure time. The Hoechst 33342 signal was recorded using a DAPI-50LP-A-000 filter set with an exposure time of 3 ms. Filter sets were from Semrock.

For analysis, ROIs were automatically generated by applying a manual threshold on the Hoechst 33342 signal, followed by utilizing the built-in particle analysis algorithm of Fiji. Data were extracted from both the green and red channels within the same ROIs. Specifically, to determine the average fluorescence intensity of cpGFP in transfected cells only, those exhibiting an mCherry fluorescence signal above a threshold of 2000 counts were considered as transfected. A more detailed description of the analysis procedure can be found in Rühl *et al.*, 2021 (35).
